# Visualizing Molecular Interactions that Determine Assembly of a Bullet-Shaped Vesicular Stomatitis Virus Particle

**DOI:** 10.1101/2022.04.07.487545

**Authors:** Simon Jenni, Joshua A. Horwitz, Louis-Marie Bloyet, Sean P.J. Whelan, Stephen C. Harrison

## Abstract

Vesicular stomatitis virus (VSV) is a negative-strand RNA virus with a non-segmented genome, closely related to rabies virus. Both have characteristic bullet-like shapes. We report the structure of intact, infectious VSV particles determined by cryogenic electron microscopy. By compensating for polymorphism among viral particles with computational classification, we obtained a reconstruction of the shaft (“trunk”) at 3.5 Å resolution, with lower resolution for the rounded tip. The ribonucleoprotein (RNP), genomic RNA complexed with nucleoprotein (N), curls into a dome-like structure with about eight gradually expanding turns before transitioning into the regular helical trunk. Two layers of matrix (M) protein link the RNP with the membrane. Radial inter-layer subunit contacts are fixed within single RNA-N-M1-M2 modules, but flexible lateral and axial interactions allow assembly of polymorphic virions. Together with published structures of recombinant N in various states, our results suggest a mechanism for membrane-coupled self-assembly of VSV and its relatives.

## INTRODUCTION

Non-segmented, negative-strand (NNS) RNA viruses of the order *Mononegavirales* include such human pathogens as Ebola virus, respiratory syncytial virus (RSV), measles virus, and rabies virus (RABV). The study of vesicular stomatitis virus (VSV), a prototypic mammalian rhabdovirus, has contributed to our fundamental understanding of the biology of NNS RNA viruses (Lyles and Rupprecht, 2007; Plemper and Lamb, 2021). The viral genome comprises a regulatory 3’ terminal leader region and 5 genes in the following order: N (nucleoprotein), P (phosphoprotein), M (matrix), G (glycoprotein) and L (large polymerase), and a 5’ terminal trailer region. All of these proteins are present in mature, infectious virions, in which the negative-strand, genomic RNA is coated with the N protein (one N per 9 ribonucleotides) to form a ribonucleoprotein (RNP) complex (Luo et al., 2020). The VSV RNP complex assembles, together with M, into a helical, bullet-shaped structure, as it buds through the plasma membrane of an infected cell (Riedel et al., 2020). Projecting from the membrane are trimeric glycoproteins (G), which mediate host cell attachment and low pH-induced membrane fusion (Kim et al., 2017; Roche et al., 2006; Roche et al., 2007). Multiple copies of L, a polyfunctional RNA polymerase, are tethered by their P cofactor to the interior of the helical RNP. Using the RNP complex as template, the L protein catalyzes transcription and replication with its RNA-dependent RNA polymerase (RdRp) activity and also caps and methylates the 5’ ends of the viral mRNA transcripts (Jenni et al., 2020; Liang et al., 2015).

Atomic structures from x-ray crystallography and cryogenic electron microscopy (cryo-EM) are available for all five VSV proteins (Ding et al., 2006; Gaudier et al., 2002; Green and Luo, 2009; Green et al., 2006; Jenni et al., 2020; Leyrat et al., 2011; Ribeiro et al., 2008; Roche et al., 2007). VSV N, expressed as a recombinant protein in bacterial cells, binds random fragments of RNA and forms ring-like assemblies (Chen et al., 2004; Green et al., 2000). A crystal structure of the predominant species, a decamer with up to 90 nucleotides of RNA, has been particularly informative for understanding the organization of the virion (Green et al., 2011; Green et al., 2006; Zhang et al., 2008). N is an elongated, two-lobed protein of ∼420 amino-acid residues; the single stranded RNA binds in a cleft between the two lobes, with nine nucleotides per N subunit. Flexible extensions from each N subunit contact the neighboring subunits along the RNA. The N-terminal arm of each subunit (residues 4–19) embraces the C-terminal lobe of the subunit 5’ to it along the genomic RNA, folding into a shallow groove; an extended loop of the C-terminal lobe (residues 342–357) contacts the C-terminal lobe of the subunit 3’ to it.

Early examination of the RNP complex in intact bullet-shaped virions by EM of negatively stained VSV particles suggested a helix of about 30 coils with an external diameter of 490 Å (Nakai and Howatson, 1968). The hemispherical tip of the bullet shape was proposed to arise from four additional turns with diminishing diameter. A more refined model was obtained from a cryo-EM reconstruction of the helical region of the virion, referred to in that work as the “trunk”, determined at about 10 Å resolution (Ge et al., 2010). Docking of the N and M protein crystal structures into the density showed that the RNP complex forms an inner, left-handed helix, surrounded by a second, outer helix of M proteins. The long axis of the N protein is approximately perpendicular to the helix axis, with its RNA-binding cleft oriented towards the tip of the virion. The 3’ end of the genomic RNA is therefore at the domed end of the particle and the 5’ end, at the blunt end of the particle.

To reach higher resolution, we recorded cryo-EM images of intact VSV virions preserved in vitreous ice and used a supervised classification approach to sort virions with different morphologies. We obtained a helical reconstruction of the trunk at 4.1 Å resolution, and a local reconstruction of the basic building module (one N and two M proteins) at 3.5 Å resolution. Our atomic model of the helical nucleocapsid shows that two concentric layers of M, named M1 and M2, coat the helically folded RNP ribbon. The assembly is held together by invariant interactions between the N, M1, and M2 subunits within the basic module, and by contacts with neighboring modules through flexible extensions. We also obtained an asymmetric reconstruction of the domed tip at about 9 Å resolution. The RNP ribbon interacts with the viral membrane at the tip of the virion, curls into about eight turns with increasing diameter, at which point the ribbon transitions into the regular helical trunk.

## RESULTS

### Cryo-EM Structure Determination of the VSV Helical Nucleocapsid

As in the published, 10 Å resolution structure (Ge et al., 2010), we did not detect regular features for the G protein, which is present in many fewer copies than N and which has no fixed position with respect to the internal assembly. We therefore concentrated entirely on the internal structures. For an initial reconstruction from trunk segments extracted from our cryo-EM images, we applied the previously reported helical symmetry (37.5 subunits per turn, 51.7 Å pitch) (Ge et al., 2010). Despite the fact that the data had been acquired with a direct electron detector, the resolution of the reconstruction was only about 8 Å, suggesting structural heterogeneity among the extracted helical segments. When we measured the length (1947 ± 76 Å) and diameter (638 ± 20 Å) of the bullet-shaped particle projections in our micrographs, we found broad distributions (Figure 1A) around the previously determined values (1960 ± 80 Å and 660 Å, respectively) (Ge et al., 2010); another study had reported a similarly broad diameter distribution for reconstituted trunk segments (Desfosses et al., 2013). Assembly of virions with longer or truncated RNA genomes could lead to particles with different length (Nakai and Howatson, 1968). Moreover, variations in the number of nucleoprotein (N) and matrix protein (M) subunits per helical coil in the trunk, and/or flattening of the particles in vitreous ice during cryo-EM specimen preparation, could explain the observed distributions in diameter. After conventional 2D classification of helical trunk segments, even the best-looking 2D class averages contained heterogeneous segments (Figure S1A). Top views of partially assembled or disrupted particles showed helical turns with different numbers of subunits (Figure S1B), and a scatter plot showed that bullet-shaped particles with a larger diameter were on average shorter (Figure S2A). These observations suggested that we would need to account for variations in the number of subunits per helical turn in order to improve the resolution of a cryo-EM reconstruction of the helical VSV nucleocapsid.

**Figure 1.**
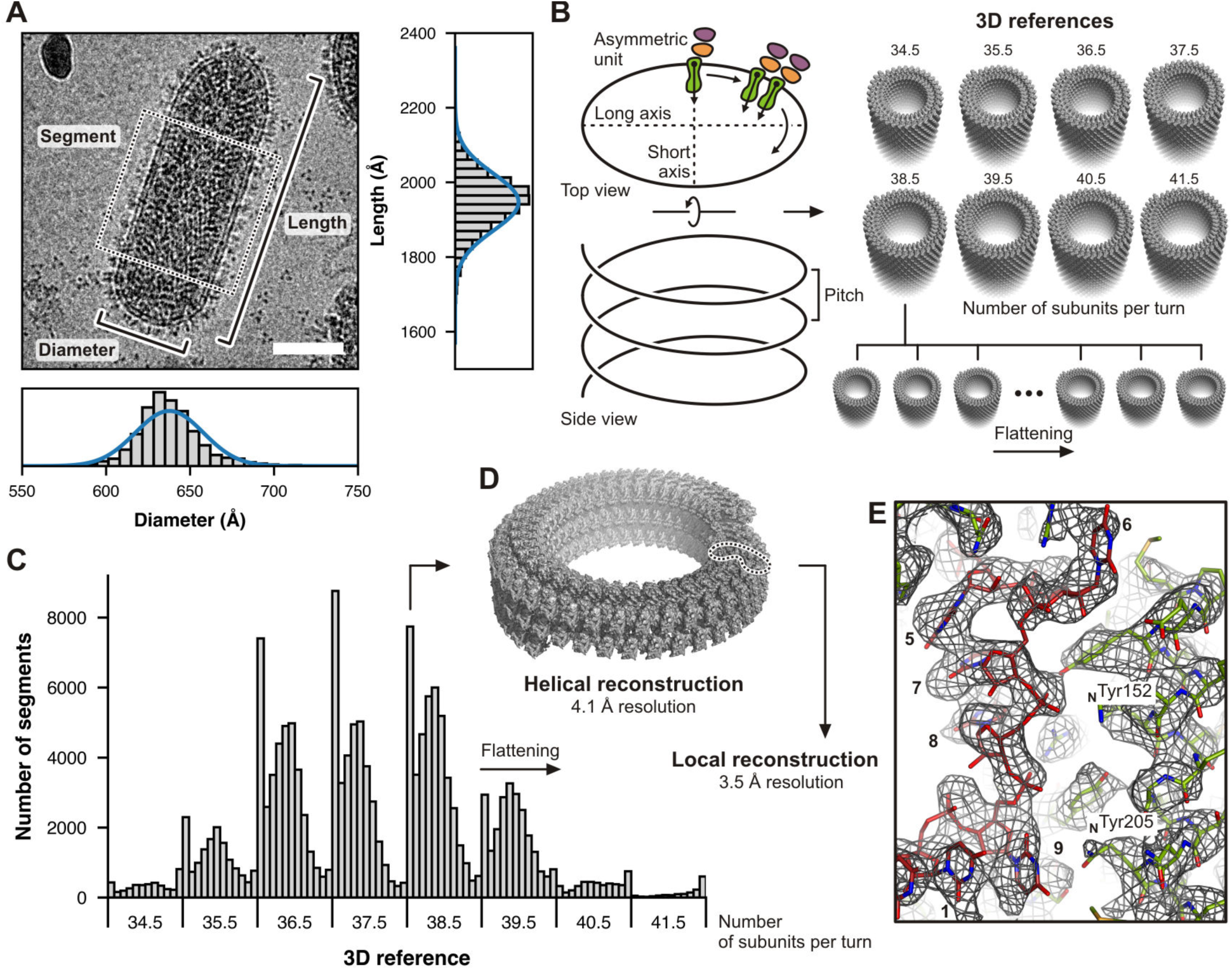
Cryo-EM Analysis of the Helical VSV Nucleocapsid. (A) Cryo-EM image of a bullet-shaped vesicular stomatitis virus (VSV) particle. The image was low-pass filtered and contrast enhanced. The scale bar corresponds to 50 nm. The horizontal and vertical histograms show the distribution of measured diameter and length, respectively, from 5763 individual particles. Measurements were taken between the middle of the lipid bilayers as indicated and possible out-of-plane tilting was ignored. The solid blue lines are normal distribution fits to the histograms with means and standard deviations of 638 ± 20 Å for the diameter and 1947 ± 76 Å for the length. The dashed box shows one segment image extracted along the VSV trunk. (B) Preparation of 3D references for supervised classification. Density of one asymmetric unit initially obtained from a low resolution reconstruction comprising one N protein (green) and two M proteins (orange and purple) was expanded to create 3D references with different numbers of subunits per turn (34.5, 35.5, 36.5, 37.5, 38.5, 39.5, 40.5, and 41.5) and different degrees of flattening. (C) Histogram of the supervised classification. Each segment image was assigned to one of the 96 3D references based on the highest alignment score. (D) Helical reconstruction obtained from sorted segments that scored best with un-flattened or only moderately flattened 3D references. Shown here is the reconstruction calculated from segments with 38.5 subunits per turn. (E) Close-up view of the cryo-EM density maps after local reconstruction of the asymmetric unit with a nominal resolution of 3.5 Å. Subparticles were aligned by alignment-by-classification and subsequent local alignment. RNA bases are numbered.

We therefore sorted helical trunk segments by supervised classification (see Method Details). To do so, we prepared 3D references with different numbers of subunits per turn and different degrees of flattening, by expanding the asymmetric unit taken from the initial low-resolution map (Figure 1B). We first globally aligned each helical segment to eight non-flattened 3D references with the number of subunits per turn, *N*, ranging from 34.5 to 41.5. Class assignment based on the highest alignment score correlated with the average measured particle length of the virions from which the segments had been extracted (Figure S2B). In a second step, we locally aligned the segments of each non-flattened class to flattened 3D references. Figure 1C shows the partitioning of segments into the final 96 classes. Helical reconstructions calculated from segments that belonged to unflattened classes showed substantially improved resolution, the best being the reconstruction obtained from the 38.5 subunits-per-turn class, which had an overall resolution of 4.1 Å as judged by Fourier shell correlation (FSC) analysis (Table S1 and Figure S3A). Note that 38.5 subunits per coil were predicted based on the diameter of the decameric N-RNA ring in the crystal structure (Green et al., 2006) and the dimension of virions observed in EM images (Nakai and Howatson, 1968).

We also calculated a local reconstruction by alignment and averaging of the asymmetric units (N subunit, 9 bound RNA nucleotides, and the two matrix protein subunits M1 and M2) from all *N* classes, yielding a nominal resolution of 3.5 Å (Figure 1E and S3B). This map was then used to build and refine a molecular model of the asymmetric unit (Table S1). We modeled the RNA nucleotides as poly-uridine, because the density in our reconstruction is the average of different sequences. The model from the local reconstruction map was then also placed into the *N* = 38.5 helical reconstruction and refined at 4.1 Å resolution (Table S1 and Figure S3C) and rigid-body fitted into the maps for the other *N* classes.

### Structure of the VSV Helical Nucleocapsid (RNA-N-M1-M2 Helix)

The asymmetric unit of the VSV helical nucleocapsid, which repeats every 9 RNA nucleotides, contains one nucleoprotein (N) subunit and two (chemically identical) matrix protein subunits (M1 and M2). Depending on the number of subunits per helical turn, the radius of the RNA strand from the center of the virion ranges from 177 Å (*N* = 34.5) to 212 Å (*N* = 41.5) (Figure 2A), in close agreement with the position of the densely stained band at a radius of 175–205 Å seen in the early EM studies of negatively stained VSV particles (Nakai and Howatson, 1968). From the length of the RNA genome (Figure 2B) and the 9-nucleotide spacing, one would expect 1,240 N proteins in a fully assembled virion, close to the value of 1,258 determined by scanning transmission electron microscopy (STEM) and biochemical analysis (Thomas et al., 1985). If one subtracts approximately 220 N subunits that form the bullet tips up to the point where the RNP ribbon transitions into a regular helix (see below), the number of coils in the helical part of the VSV nucleocapsid is 24.6–29.6, with 26.5 in case of the predominant class with *N* = 38.5.

**Figure 2.**
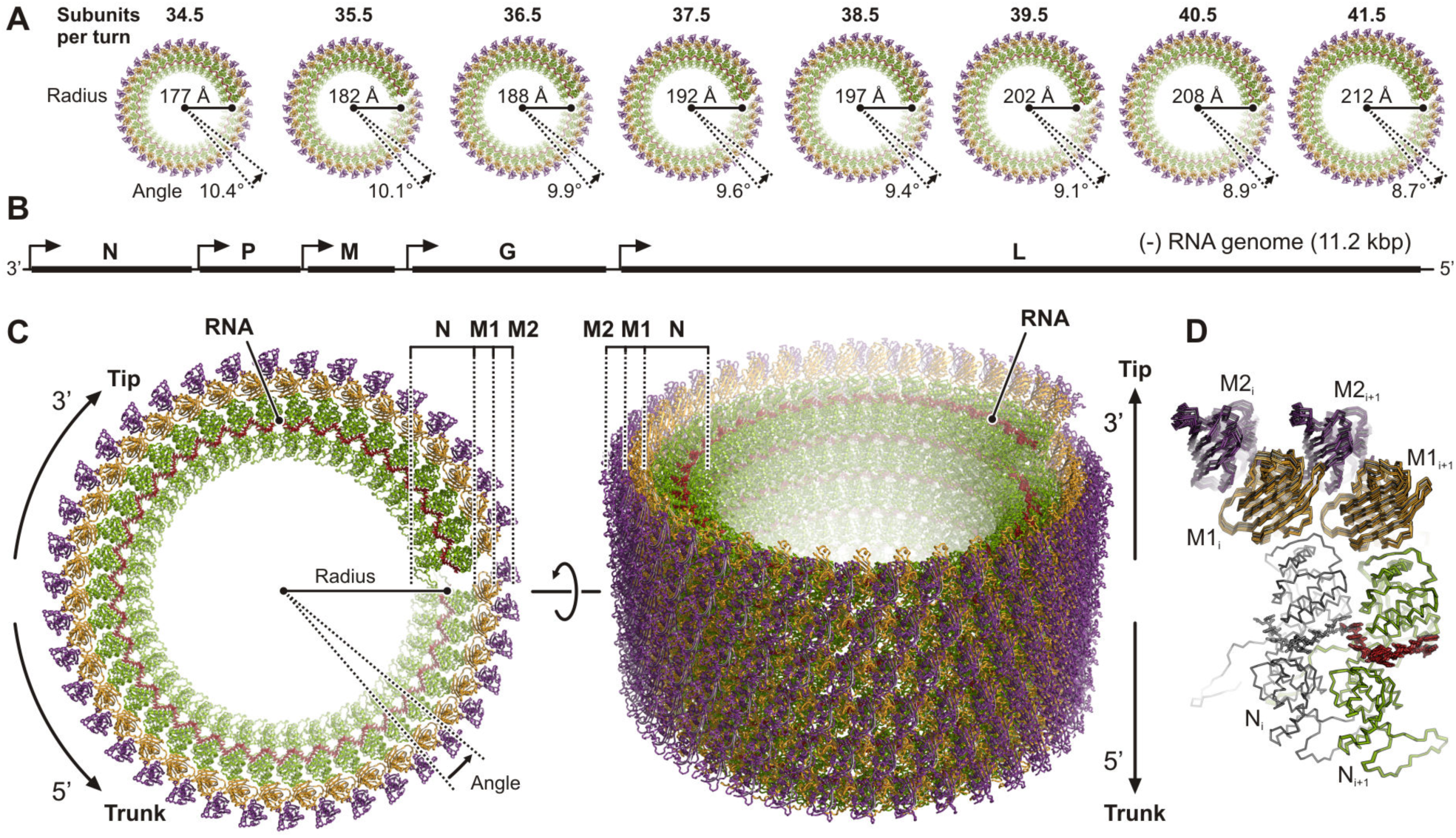
Structure of the Helical VSV Nucleocapsid. (A) Top views of the helical nucleocapsids from virions with different numbers of subunits per turn. The nucleoprotein (N) is colored green, the RNA red, and the two matrix proteins (M1 and M2) are colored orange and purple, respectively. (B) Schematic organization of the VSV negative-strand genomic RNA. Open reading frames encoding the five viral proteins are indicated by solid black lines. Transcription start sites are indicated by arrows. The 3’ end of the single-stranded RNA terminates at the tip of the virion, the 5’ at the base of the trunk. (C) Structure of the helical nucleocapsid with *N* = 38.5 in top (left) and side (right) view. Molecules are colored as in (A). The three protein layers are labeled N (nucleoprotein), M1 (inner layer matrix protein), and M2 (outer layer matrix protein). (D) Structural comparison of the subunit arrangement in nucleocapsids with different *N*. Two adjacent asymmetric units (RNA-N-M1-M2) of each class are shown in ribbon representation and are superimposed one N protein (colored in gray).

The VSV helical trunk has three concentric protein layers (Figure 2C). The innermost layer is the RNA-N protein (RNP) complex, which coils into a left-handed helix with a polarity such that the 3’ end of the (-)-strand genomic RNA is at the tip of the bullet-shaped virions and the 5’ end at the blunt end of the trunk. Two layers of matrix protein (M) surround the RNP, with the same helical symmetry. In the previous trunk structure, only a single matrix protein layer was placed into the density map, presumably because of the limited resolution of the reconstruction (Ge et al., 2010). A local resolution estimate for the *N* = 38.5 helical reconstruction (Figure S3D) and analysis of the refined temperature factors in our model (Figure S3E) show that the outer matrix protein (M2) layer is less well defined than the inner matrix protein (M1) layer. We explain this observation by partial occupancy of M2-layer subunits in the outer layer, consistent with the reported stoichiometry of only 1.5 M proteins per N protein in virions (Thomas et al., 1985). The best-ordered parts of the trunk were the M1 layer and the N-terminal domain (N lobe) of the N protein (Figure S3D and E).

As evident from the superposition of two adjacent asymmetric units on a single common nucleoprotein subunit (Figure 2D), the molecular contacts that govern coiling of the RNP ribbon into a helix are not fundamentally altered when the number of subunits per helical turn changes from 34.5 to 41.5 between different virions. Our classification analysis suggests a Gaussian distribution with an energy minimum for *N* = 37.5 or 38.5 (Figure 1C).

The structures of the N and M subunits in the assembled nucleocapsid determined here (Figure 3) are generally as observed in previous crystal structures of the individual proteins (Gaudier et al., 2002; Graham et al., 2008; Green and Luo, 2009; Green et al., 2011; Green et al., 2006; Hanke et al., 2017; Leyrat et al., 2011; Zhang et al., 2008) (Figure S4A–E). Figure S4F shows per-residue Cα distances after superposition of our N, M1 and M2 structures onto PDB-ID 2GIC (N) and PDB-ID 1LG7 (M), respectively. Most apparent are the different conformations of the N-terminal arm and the extended loop as they pass into contact with the neighboring subunit. While the conformation of residues in those segments that are directly engaged in binding the adjacent subunits are the same (Figure 3A and B), their linkages are flexible and allow the protein to adopt a more tightly curved arrangement of N in crystallized RNA-N decamers than in the trunk of assembled virions (Figure S4C). We also found conformational differences in two N protein loops (residues 111–133 and 166–181, respectively) (Figure S4C).

**Figure 3.**
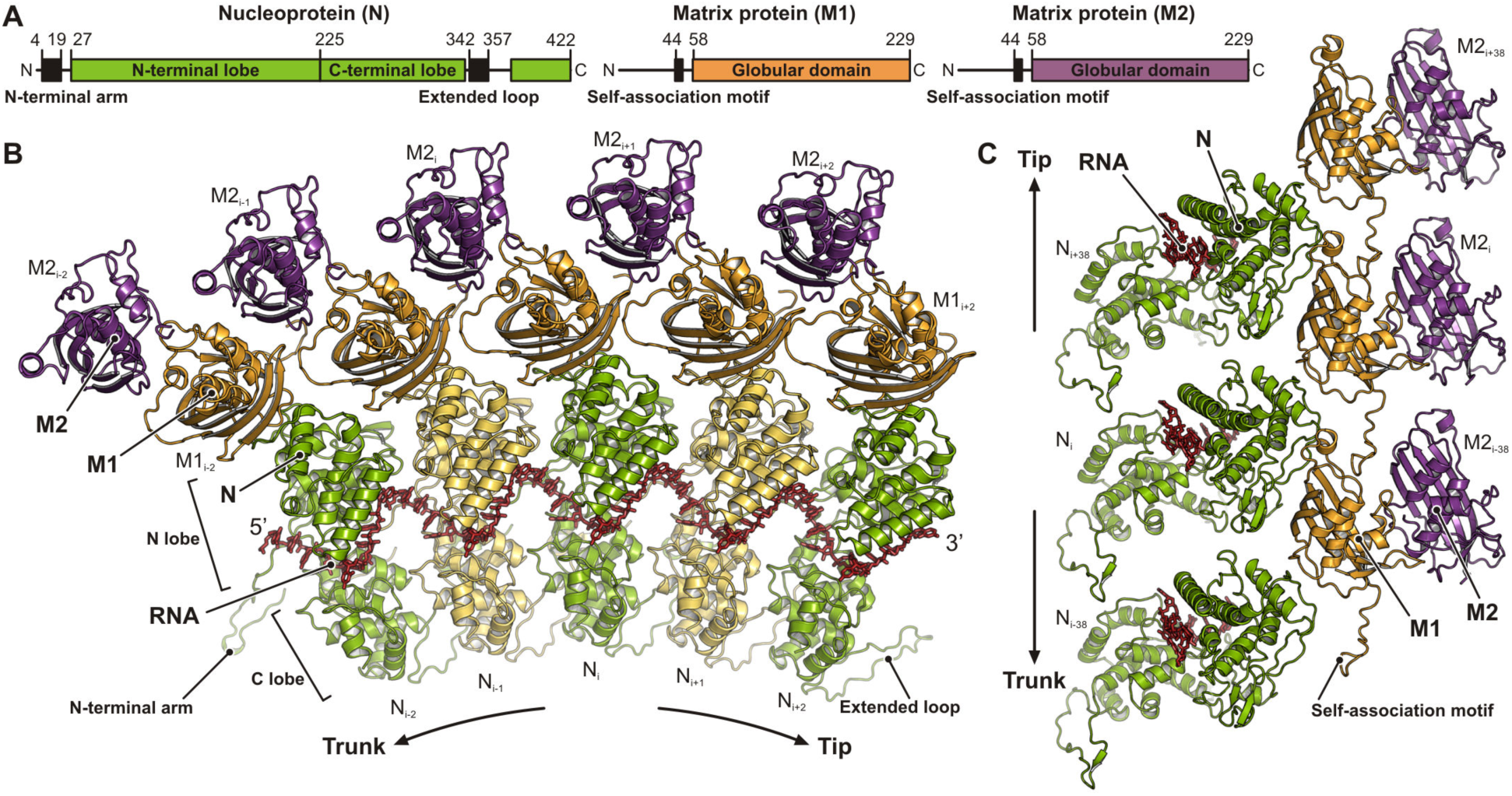
Structure of the RNA and Protein Subunits within the VSV Nucleocapsid. (A) Linear domain organization of the nucleoprotein (N, green) and matrix proteins (M1 and M2, orange and purple, respectively). Amino acid numbers indicate domain boundaries. Globular domains are shown in color. Sequences engaged in inter-subunit contacts are shown as black boxes. (B) Top view of five consecutive asymmetric units along the RNP ribbon. The RNA is shown in red stick representation. Nucleoproteins (N) are colored alternating in green and yellow. The matrix protein of the inner layer (M1) is colored in orange, the one of the outer layer (M2) in purple. Subunits are labeled with indices that increment towards the virion tip, where the 3’ end of the RNA is located, and decrement towards the base of the trunk, where the 5’ end of the RNA is located. (C) Side view of the asymmetric units from three turns of the RNP ribbon. RNA and subunits are colored as in (B). The tip of the virion would be at the top, the base of the trunk at the bottom. Indices of the individual subunits are shown.

The RNA strand lies in a cavity between the N and C lobes of the nucleoprotein (Figure 3B and C). A different conformation of nucleotide 9, either flipped out or base-stacked on nucleotide 1, was observed in various crystal structures (Green et al., 2011) (Figure S4D) and also when compared to the rabies virus (RABV) RNP structure (Albertini et al., 2006). In the virion structure here, nucleotide 9 base stacks with nucleotide 1 and is part of a first quasi-helical stretch (Green et al., 2006) together with nucleotides 1–4. The second quasi-helical stretch comprises base-stacked nucleotides 5, 7, and 8. Nucleotide 6 bulges out, separating the two stretches, and its base is not well resolved in our density map. We mapped the N-protein residues that contact the RNA in our structure onto a multiple-sequence alignment that includes sequences from other viruses of the *Rhabdoviridae* family (Data S1). Conserved residues _N_Arg143, _N_Lys206, _N_Arg214, _N_Lys286, _N_Arg408, all ordered in our reconstruction with well-resolved side chain density, coordinate the phosphate groups of the RNA backbone by salt bridges. _N_Arg312 probably also contributes a salt bridge, but its side chain is less well resolved. Conserved _N_Tyr152 hydrogen- bonds to the phosphate group of nucleotide 8, and stacks with its aromatic side chain onto the sugar ring of nucleotide 6. The side chains of _N_Arg408 and _N_Arg317 stack on the bases of nucleotides 6 and 4, respectively. Other conserved residues in the binding cleft establish shape complementarity and engage in hydrophobic contacts with the RNA strand. These contacts do not depend on the identity of the base at each nucleotide position and therefore allow the N protein to bind any 9-base sequence in the RNA genome.

### Subunit Interactions in the VSV Nucleocapsid

To describe the subunit interactions within the helical nucleocapsid, we classified the interfaces according to whether they are within a layer (lateral and axial intra-layer contacts for the N, M1 and M2 layers) or between layers (radial inter-layer contacts). Table 1 summarizes the observed interfaces in the *N* = 38.5 helical structure together with the calculated interface areas (IA), or buried accessible surface areas, for each of the contacts. Interacting residues are labeled in the multiple sequence alignments in Data S1–3.

**Table 1.** Subunit Interactions in the VSV Nucleocapsid.

| **Intra-layer N contacts** |  |  | **IA<sup>a</sup> (Å<sup>2</sup>)** |
| --- | --- | --- | --- |
| Lateral intra-layer contacts |  |  |  |
| Interface 1 | Single stranded RNA bound to N subunit |  | 1055 |
| Interface 2 | N<sub>i</sub> N-terminal arm interacts with N<sub>i-1</sub> C-terminal lobe and with N<sub>i-2</sub> extended loop |  | 793, 354 |
| Interface 3 | N<sub>i</sub> extended loop interacts with N<sub>i+1</sub> C-terminal lobe and with N<sub>i+2</sub> N-terminal arm |  | 683, 354 |
| Interface 4 | N<sub>i</sub> N-terminal lobe interacts with N<sub>i+1</sub> N-terminal lobe |  | 442 |
| Interface 5 | N<sub>i</sub> C-terminal lobe interacts with N<sub>i+1</sub> C-terminal lobe |  | 207 |
| Axial intra-layer contacts |  |  |  |
| Interface A | N<sub>i</sub> subunit interacts with N<sub>i+39</sub> subunit |  | 94 |
| Interface B | N<sub>i</sub> subunit interacts with N<sub>i+38</sub> subunit |  | 18 |
| **Intra-layer M1 contacts** |  |  | **IA (Å<sup>2</sup>)** |
| Lateral intra-layer contacts |  |  |  |
| Interface 6 | M1<sub>i</sub> globular domain interacts with M1<sub>i+1</sub> globular domain |  | 65 |
| Axial intra-layer contacts |  |  |  |
| Interface C | M1<sub>i</sub> self-association motif interacts with M1<sub>i-38</sub> globular domain |  | 400 |
| Interface D | M1<sub>i</sub> globular domain interacts with M1<sub>i+39</sub> globular domain |  | 78 |
| **Intra-layer M2 contacts** |  |  | **IA (Å<sup>2</sup>)** |
| Axial intra-layer contacts |  |  |  |
| Interface E | M2<sub>i</sub> globular domain interacts with M2<sub>i+38</sub> globular domain |  | 49 |
| **Inter-layer N-M1 contacts** |  |  | **IA (Å<sup>2</sup>)** |
| Radial inter-layer contacts |  |  |  |
| Interface i | N<sub>i</sub> C-terminal lobe interacts with M1<sub>i</sub> globular domain |  | 345 |
| Interface ii | N<sub>i</sub> C-terminal lobe interacts with M1<sub>i-38</sub> globular domain |  | 310 |
| Interface iii | N<sub>i</sub> C-terminal lobe interacts with M1<sub>i+1</sub> globular domain |  | 117 |
| **Inter-layer M1-M2 contacts** |  |  | **IA (Å<sup>2</sup>)** |
| Radial inter-layer contacts |  |  |  |
| Interface iv | M1<sub>i</sub> globular domain interacts with M2<sub>i</sub> globular domain |  | 600 |
| Interface v | M1<sub>i</sub> globular domain interacts with M2<sub>i+1</sub> globular domain |  | 430 |

<sup>a</sup> IA: interface area (buried accessible surface area upon interface formation).

#### Intra-layer contacts

The N protein clamps onto the RNA strand (interface 1) and forms extensive contacts with adjacent N subunits thought its N-terminal arm and extended loop (Figure 4A, bottom). The segments of these extensions that bind the C lobes of the preceding (interface 2) and following (interface 3) N subunits, respectively, are essentially as observed in the crystal structures. The N-terminal arm of an N subunit folds through a groove in the subunit 5’ to it along the RNA; the extended loop folds against the blunt-end facing surface of the subunit 3’ to it and also interacts with the N-terminal arm of the next N subunit in the 3’ direction. These elaborate imbrications would tie together the N-protein ribbon even without the continuity of the RNA polynucleotide chain, and we have suggested elsewhere that they indeed do so when the RNA passes into the polymerase active site, retaining continuity of the RNP upstream and downstream of the “transcription bubble”. The tethers are flexible as they pass from their subunit of origin into the neighboring subunit, allowing variability of overall radius of the RNP helix, as seen in the widening turns at the tip and in the distribution of trunk diameters, while maintaining a constant 9-nucleotide spacing along the RNA strand. When the RNP complex coils into the helical nucleocapsid, there are additional lateral contacts between N-protein neighbors (Figure 4A bottom). Between adjacent N-lobes, salt-bridges form between _N_Glu169 and _N_Arg179 and between _N_Asp85 and _N_Lys34 (interface 4). C-lobe contacts (interface 5), closer to the helix axis, are much less extensive. Axial contacts in the N layer (interfaces A and B) are likewise tenuous, where N_i_ fits loosely into the shallow groove between N_i+38_ and N_i+39_ (Figure 4A, top).

**Figure 4.**
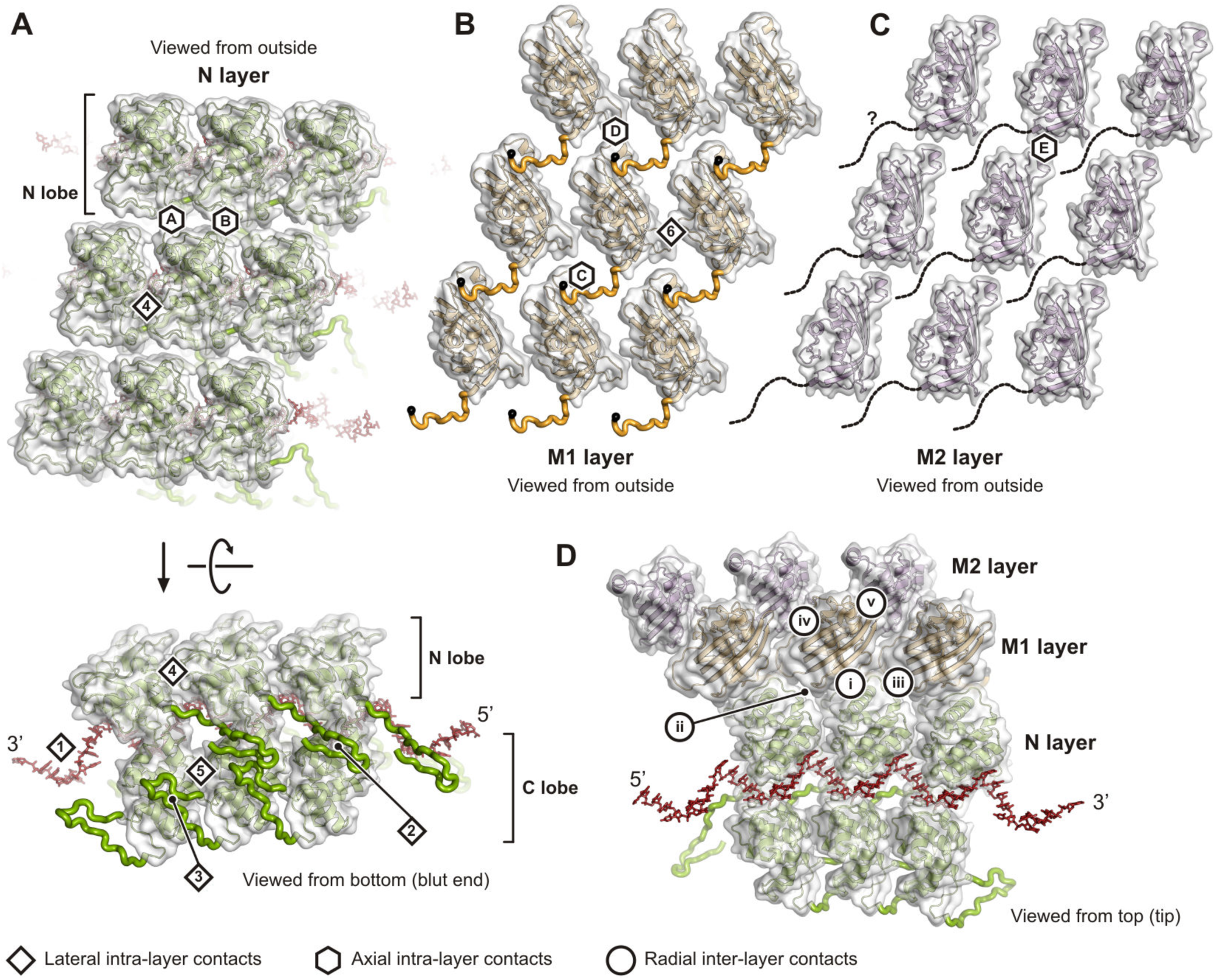
Inter-Subunit Contacts. RNA is shown in red stick representation. Globular domains are shown with transparent surface, extensions as thick ribbon. Lateral intra-layer contacts are labeled with diamonds and Arabic numerals, axial intra-layer contacts with hexagons and letters, and radial inter-layer contacts with circles and Roman numerals. See also Table 1. (A) Inter-subunit contacts within the N protein layer. Top, side view of the N layer from the outside. Bottom, bottom view of the N layer. (B) Inter-subunit contacts within the M1 protein layer in side view from the outside. (C) Inter-subunit contacts within the M2 protein layer in side view from the outside. (D) Inter-subunit contacts between the N, M1 and M2 layers.

In the only substantial intra-layer contact in the M layers, the “self-association motif” (residues 44–52) (Graham et al., 2008) creates a chain of interactions within the M1 layer, and possibly within the M2 layer (Figure 3A, C and Figure 4B, C). The motif from M1 subunit i interacts with a pocket defined by residues 78–84 and 114–125 in M1 subunit i+38 (interface C), as predicted from lattice contacts in crystals of VSV M (Graham et al., 2008). Poor M2-layer resolution prevented us from determining whether the same axial contact is also present there; the positions and orientations of M2 subunits would allow the contact to be present. Other intra-layer contacts (interfaces 6, D and E) are very slight and unlikely to contribute to overall stability (Table 1).

#### Inter-layer contacts

Each N protein contacts three M1 subunits (Figure 4D and Table 1). The first of these radial inter-layer contacts, interface i, is the most extended, and the one that is also present in the tip of the VSV bullet (see below). The contact is well resolved in our local reconstruction and involves N protein residues 116–125 from a long loop that wraps around the N lobe domain and binds two M1 subunit loops, residues 94–101 and 149–152. The M1 residues of this interface are conserved (Figure S5 and Data S2). Stabilizing interactions include hydrogen bonds between the main chain of the N loop 116–125 with the M1 loop 94–101. We also observe hydrophobic contacts between residues _N_Val121 and _M1_Met98 and _N_Val116-_N_Leu117-_N_Pro118 and _M1_Pro149-_M1_Pro150. The second contact of N with an M subunit, interface ii, is with M1_i-38_. Salt bridges are formed between _N_Lys78 and _M1_Glu130, and between _N_Asp100 and _M1_Arg159. Most other residue contacts in this interface are also polar. The third, minor contact, interface iii, is between the N protein C-terminal lobe and the M1_i+1_ globular domain.

Each M1 protein contacts two M2 subunits (Figure 4D and Table 1). Interface iv is an extended contact with the M2_i_ subunit. _M1_Asp191 binds _M2_Arg102. The M2 C terminus with its _M2_Phe228 accommodates in a hydrophobic cleft of M1 where it interacts with _M1_Trp220. Additional hydrophobic contacts are between _M1_Pro187 and _M2_Met94-_M2_Ile96. Interface v is a contact with the M2_i+1_ subunit.

The buried accessible surface areas for each interface (Table 1) and residue conservation analysis (Figure S5) support the notion that the VSV nucleocapsid is held together primarily by lateral interactions in the N layer (interfaces 1–3), axial interactions in the M1 layer (interface C), and radial interactions between the layers (interfaces i and iv). Indeed, surface residues of those interfaces generally show the highest conservation, and deletion of resides in the N-terminal arm or extended loop both result in loss of N oligomerization (Zhang et al., 2008). Proteolytic cleavage of the M protein likewise prevents self-association in vitro (Gaudier et al., 2002).

### Structure of the VSV Tip

With a stepwise particle alignment and reconstruction protocol, which we describe in detail in the Methods section and Figure S6, we obtained an asymmetric reconstruction of the rounded VSV tip (Figure 5A). The reconstruction comprising the first ten turns of the RNP ribbon had a nominal resolution of about 9 Å and allowed a rigid body-fit of the structures of individual modules (Figure S7). The density along the RNP ribbon showed that binding of M1 to N (interface i) within each module in the tip is identical to the corresponding interaction in the modules of the trunk. The same appears to be true for binding of M2 to M1 (interface iv), but partial occupancy of the M2 subunit resulted in weak density for the M2 layer generally.

**Figure 5.**
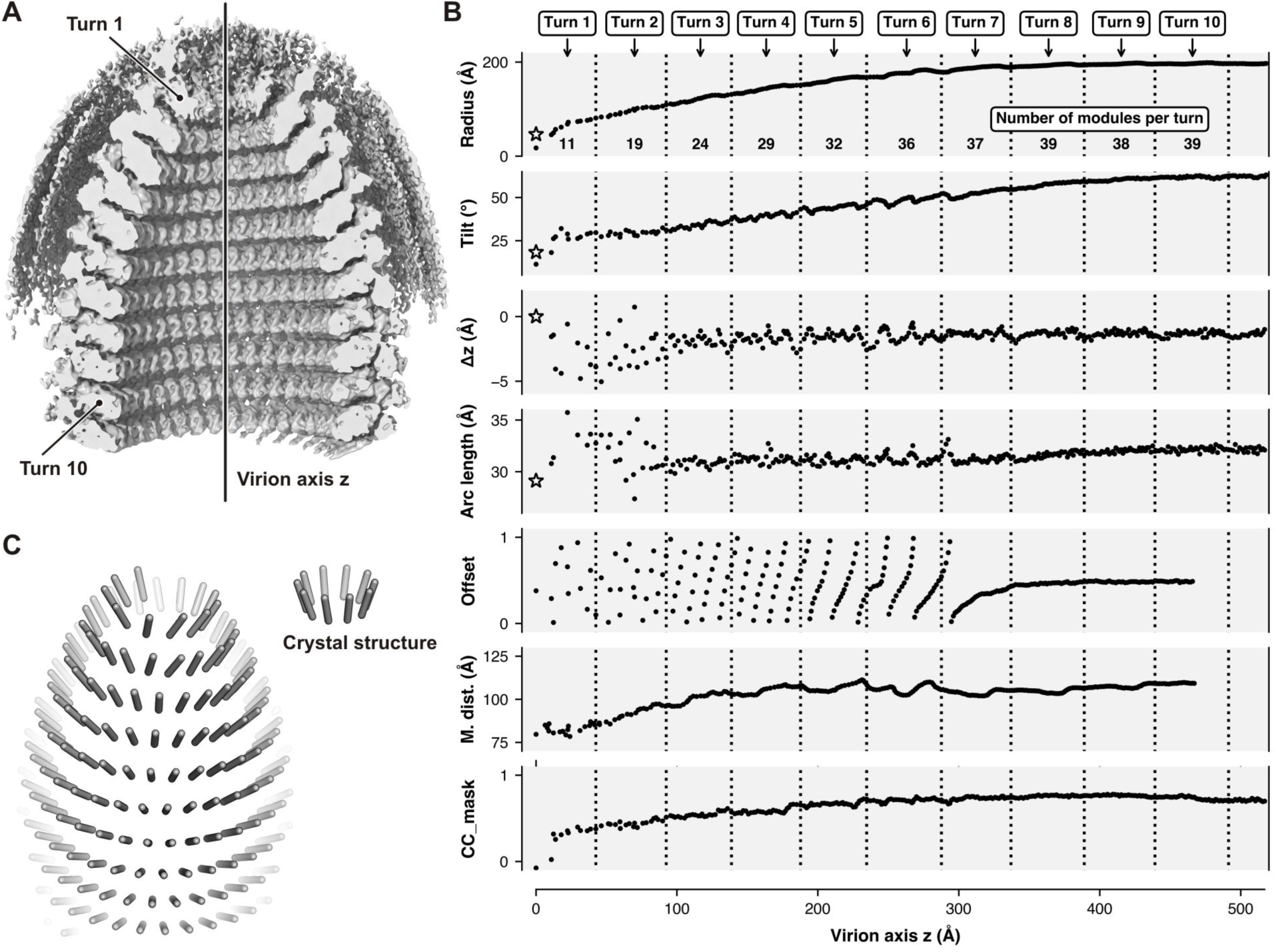
Structure of the VSV Tip. (A) Asymmetric reconstruction of the VSV tip. The density was low pass filtered at 8.8 Å resolution and sharpened with by applying a *B* factor of -100 Å^2^. To allow inside view, the density is partially cut. (B) Analysis of individual modules after rigid body fitting into the map of the VSV tip reconstruction. The horizonal axis is the distance from the top of the tip along the virion axis. Each dot corresponds to one module. Turns are delineated on top of the plot. Asterisks in the plots are values calculated from the crystal structure of the decameric N-RNA complex (Green et al., 2006). Radius: distance between the virion axis and the center of gravity of the nine RNA bases. Tilt: tilt of the N protein relative to the helical axis. Δz: position of the module along the virion axis (RNA center of gravity). Arc length: lateral distance between adjacent modules (RNA center of gravity). Offset: Relative offset of the module with respect to the position of the modules in next turn (dihedral angels, RNA center of gravity). M. dist.: distance between the module (RNA center of gravity) and the middle of the lipid bilayer. CC_mask: correlation coefficient between the rigid-body fitted module (N and M1 subunits) and the map of the tip reconstruction. (C) Schematic representation of the N subunit arrangement in the VSV tip. The longitudinal axis of each N subunit is shown (left) and compared to the N arrangement in the decameric crystal structure (right).

The first turn of the RNP ribbon at the tip of the virus, where the 3’ end of the genomic RNA is located, comprises about 11 modules. The number of modules then increases in each subsequent turn, up to turn eight, where we counted 39 modules, which corresponds to the number of modules per turn in the regular helical structure of the nucleocapsid trunk (Figure 5B). A plot of the radial distance between the nine RNA bases (center of gravity) of each fitted module and the virion axis shows that the radii increase gradually from one module to the next (Figure 5B). Concomitant with this radial expansion, the tilt of modules steadily increases for an accumulated change of about 40° (Figure 5C). The relative shift along the virion axis and the arc length between adjacent N subunits are both approximately constant. The constant lateral distance between neighboring N proteins allows the 9 nucleotide-register of RNA binding (interface 1) as well as interaction of the N-terminal arm (interface 2) and the extended loop (interface 3) to be maintained, independent of the curvature of the turns. The axial interfaces between subunits that we observed after the tip-to-trunk transition (Table 1) are however not conserved in coils of the tip. The difference is evident from the offset plot (Figure 5B). In the trunk the offset is constant at 0.5, meaning that each N subunit sits exactly between two N subunits of the next turn. In the tip, however, the offset oscillates between 0.0 and 1.0, even within a single turn. The RNP ribbon in the tip essentially “slides” on the preceding turns without establishing invariant axial binding interfaces. We also measured the distance between the inner membrane leaflet and the nucleocapsid, which shows that the spacing is approximately 20 Å shorter at the tip than in the trunk (Figure 5B).

### Binding of the Phosphoprotein P to the Nucleocapsid

Mature VSV virions contain approximately 50 copies of the L protein (Thomas et al., 1985), which are incorporated into virions by its phosphoprotein (P) cofactor (Figure S8A). From published work, we know the structures in isolation of the P_L_ segment bound to the L protein (Jenni et al., 2020) and the P_CTD_ bound to the N protein (Green and Luo, 2009). We found density of the P_CTD_ domain *in situ*, bound to the N protein in the interior of the virus, when we low-pass filtered and low-level contoured maps of our trunk reconstructions (Figure S8B). Attempts to classify sub-particles for modules with bound P_CTD_ were not successful, likely because of the size of the P_CTD_ domain (8 kDa) and the relatively low occupancy. Nonetheless, the observation of P_CTD_ at low resolution in a conformation essentially as in the crystal structure (Green and Luo, 2009) further supports the current model of how P tethers L to the RNP for packaging into the virion (Figure 6).

**Figure 6.**
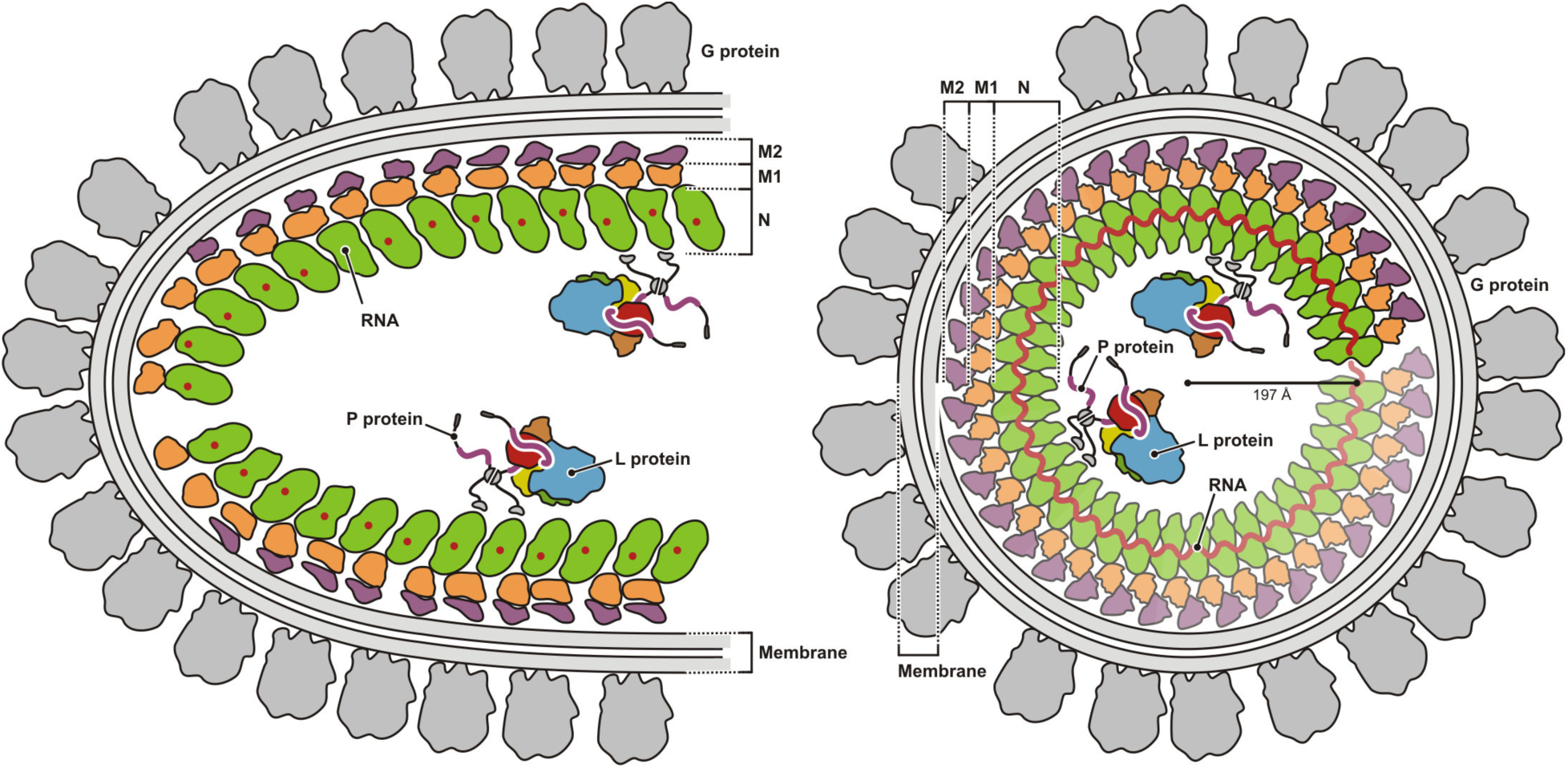
Model of VSV. Schematic diagram of VSV. A longitudinal section of the VSV tip is shown on the left, a cross section of the helical trunk on the right. All viral proteins are drawn to approximate scale.

### Membrane Association of M and G

The N-terminal extensions of M subunits in both layers project toward the membrane (Figure S8C, D). A segment containing eight lysines (“basic N terminus”, residues 5–19) crosslinks efficiently to photoactivatable membrane probes in VSV (Lenard and Vanderoef, 1990), and 24 residues then intervene between the C-terminus of this cross-linkable segment and the N-terminus of the ordered self-association motif (residue 44). In our reconstruction of the helical trunk, a gap of about 30 Å separates the M2 layer of the nucleocapsid from the inward-facing headgroups of the viral membrane (Figure S8D). If the M2-layer self-association motif were to dock to an axially shifted subunit, as in layer M1, 24 residues could readily bridge the gap between residue 44 and bilayer, and even the corresponding segment from, M1 could do so, as the distance (approximately 50 Å) is only slightly greater than 2 Å per residue. We do not see ordered density for the G protein, consistent with biochemical evidence that the cytoplasmic tail of G does not interact strongly with the matrix protein (Lyles, 2013) and with observations that infectious chimeric viruses can be engineered in which other viral fusion proteins substitute for VSV G (Roberts and Rose, 1999).

## DISCUSSION

High-resolution cryo-EM reconstructions depend on averaging images of a set of homogenous particles (in composition and or conformation) after determining their projection angles and shifts. Classification of heterogenous images can be particularly challenging if the individual particles have a high degree of local symmetry (tip structure here) or have different, but related strict helical symmetries (trunk structures here). Our supervised classification protocol partitioned the virus images into classes with different helical symmetries (number of subunits per turn) and subsequent locally focused alignment of sub-particles yielded a reconstruction of the basic trunk-building module at 3.5 Å resolution (Figure 1). For the tip reconstruction (Figure 5) we used a stepwise alignment protocol (Figure S6). The barely sub-nm resolution suggests that there is considerable heterogeneity among individual tip structures; electron cryotomographic (cryo-ET) methods might be required to understand their structural variation.

The overall resolution of the reconstruction we have obtained, and the even higher resolution of an N-RNA-M1-M2 module, has allowed us to integrate published observations into a likely sequence of events during VSV budding at the plasma membrane of an infected cell. Comparison of the initial turn of the RNP helix with the crystal structure of an RNP decamer (Green et al., 2006) shows that the lateral interactions and tilt of the subunits with respect to the axis are very similar (Figure 5C). Recombinant rabies virus N forms a closely related undecameric ring (Albertini et al., 2006). When expressed in insect cells, N binds longer RNA molecules and forms undulating ribbons with varying local curvature (Desfosses et al., 2013). At very low ionic strength and neutral pH, these ribbons condense locally into strings of unidirectional, tip-like conical structures that expand in 5 turns from a tight ring at their apex to a diameter of about 390 Å, and at pH 5, the RNP forms bullet-shaped structures very similar to the RNP in virions, but with a somewhat narrower diameter, corresponding to 32–34 subunits per turn (Desfosses et al., 2013). These observations show that N alone can determine the size and shape of the particle and that M, which appears to modulate slightly the most stable diameter for a continuous helical assembly in the trunk of a particle, couples assembly of the RNP bullet to the membrane though which it buds.

Comparing the structure of the first 8 turns of the RNP helix at the tip of the virion with published structures of N and RNPs in various forms as just described, we can propose the following picture for VSV assembly at the membrane. Budding begins at the domed end, with the 3’-end of the RNP. The similarity of the crystallized RNA ring and the first turn of the RNP in virions suggests that the ring is a good approximation to the structure that initiates virion assembly. When bound on an RNA molecule longer than 90 nucleotides, a decameric ring cannot close on itself and must instead dislocate into a lockwasher and propagate as a helix. Stacking of N subunits in successive turns, even if relatively non-specific, and the steep tilt of the subunits will then cause the second turn to have a larger diameter than the first, and similarly in increasingly wide gyres until a stable axial stacking occurs after about 8 turns, with 38.5 N subunits per turn. The energy required to curve the membrane will also favor successively greater diameters as the tip propagates, until a balance between membrane deformation and stable, half-offset axial packing occurs at the tip-to-trunk transition.

The orientation with which N binds RNA requires that the 3’ end of the genome be at the domed end, with an exposed extended loop of the C lobe of the first N protein. Initiation of assembly at the membrane might simply be a linkage through M of the subunits in the first, 10–11 subunit turn of the RNP with the inner leaflet of the membrane lipid bilayer. Alternatively, a specialized organization of the 3’-end of the RNP might create a more specific initiation complex. The resolution of the tip structure at that position does not allow a firm conclusion. One speculative possibility is that the RNA 3’ end associates with the template entry channel of a molecule of the viral RdRp L protein, just as it does to impart specificity of viral RNA plus-strand packaging in the dsRNA viruses, with one polymerase per genome segment (Ding et al., 2019; Jenni et al., 2019; Pan et al., 2021). Such an arrangement could be coupled to M-protein suppression of L catalyzed polymerization (Clinton et al., 1978). Influenza virus assembly also includes a 3’-end localized RdRp molecule on each of the eight genomic RNAs (Te Velthuis et al., 2021).

Detecting and visualizing the M2 layer and building a well-positioned model for both M1 and M2 are important consequences of the resolution of our reconstruction. Previous work had assumed that the “extra” M (present in total about 1.5 times the level of N) decorated the inner surface of the helical RNP assembly, as previously suggested in the “cigar” model of virus assembly in which the N-RNA wraps around a core of M (Barge et al., 1993). The presence of two layers of M around the N-RNA now provides an explanation for the remaining M in the virion. A key interaction between successive turns of the helical ramp is docking of the M1-layer self-association motif from subunit i into the pocket that receives it on subunit i+38. Although the short segment connecting the motif itself with the globular region of M may be flexible, a loop from subunits in the M2 layer bears against it, potentially reinforcing its conformation helping to establishing the half-staggered offset of successive turns. In the tip, where the offset varies, the M2 layer appears to be absent.

The M2 layer is only half occupied. Our current model does not show whether its distribution is systematic or random. The self-association motif could establish some degree of longitudinal coherence, so that variable-length strips of motif-linked, M2-layer subunits would decorate the outside of the M1 layer. A cryo-ET structure of the RABV trunk shows only a single M layer (Riedel et al., 2019) and further studies should clarify functional roles of variations in the nucleocapsid organization among virions of the *Rhabdoviridae* family.

## METHOD DETAILS

### Virus Production and Purification

VSV Indiana serotype was grown on BSR-T7 cells seeded in DMEM - 10% FBS, and infected one day later at a multiplicity of infection (MOI) of 3 for 1 h. The virus suspension was then replaced by DMEM - 2% FBS, and cell supernatant was harvested 20 h post-infection. The supernatant was first clarified by centrifugation at 750 x g for 5 min, and concentrated through a 15% (w/v) sucrose cushion in NTE buffer (10 mM Tris-HCl pH 7.4, 100 mM NaCl, 1 mM EDTA) at 110,000 x g for 2 h at 4 °C. Pellets were resuspended overnight at 4 °C in NTE buffer, put on top of a linear 15-45% (w/v) sucrose gradient prepared in NTE buffer, and centrifuged at 200,000 x g for 3 h at 4 °C. Band corresponding to virus was collected by side-puncture of the tube in a volume of 0.5 mL, and dialyzed overnight at 4 °C against 1 L of NTE buffer using a Slide-a-Lyzer, 0.1–0.5 mL, 10 kDa cut off, cassette.

### Specimen Preparation and Cryo-EM Data Collection

We immobilized VSV particles in vitreous ice using a CP3 cryoplunger. 3 μl of band-purified virus was applied to copper mesh holey carbon grids (C-flat 1.2-micron diameter holes with 1.3- micron spacing and a support thickness of ∼40 nm) and blotted for 3 s in a chamber maintained at 90% humidity before plunge-freezing into liquid ethane at -171 °C. Images of VSV particles were collected on a Titan Krios G3i microscope (Thermo Fisher Scientific) operated at 300 kV, and equipped with a pre-camera energy filter (Gatan Image Filter) and K3 Summit direct detector (Gatan). The nominal magnification was 105 kx, corresponding to a calibrated physical pixel size of 0.85 Å. Dose-fractionated movies were recorded in counting mode with 0.05 s per frame for a total of 2.5 s (50 frames, total dose of 60 electrons per Å^2^). Using SerialEM v3.7 (Mastronarde, 2005), we recorded a total of 27 movies for each stage position (nanoprobe mode with an illuminated area of ∼1 μm, 3 exposures per hole from a total of 9 holes).

### Cryo-EM Data Processing

#### Particle picking and movie processing

Movie frames were aligned and summed using MotionCor2 (5×5 patch alignment) (Zheng et al., 2017). In four times binned and low-pass filtered sums (18,353 micrographs) we manually marked 12,348 VSV trunks with e2helixboxer.py from EMAN2 (Bell et al., 2016). We next estimated magnification distortion from power spectra of trunks extracted from the original images, as previously described (Jenni and Harrison, 2018), applied the fitted values (distortion angle = 109.96, major scale = 1.0026, minor scale = 0.9974) with mag_distortion_correct (Grant and Grigorieff, 2015) to the original movie stacks, and used again MotionCor2 through the RELION (Scheres, 2012) wrapper relion_run_motioncorr to obtain sums from the magnification corrected movie stacks.

#### Particle extraction and CTF determination

Determination of contrast transfer function (CTF) parameters, extraction of helical trunk segments, and particle stack preparation involved the following steps: (i) Manually picked trunk coordinates from EMAN2 were converted to RELION star file format. (ii) Using relion_preprocess, we extracted helical segments based on the trunk coordinates (box size = 1200×1200 pixels, helical_nr_asu = 37.5, helical_rise = 1.38 Å). Background radius was set to 482 pixels and the helical outer diameter to 820 Å. Segment images were normalized and contrast-inverted. (iii) We used Gctf (Zhang, 2016) to determine CTF parameters from the total-summed images and refinement for values at individual segment coordinates. (iv) Unbinned and binned particles stacks were created from the extracted segment images. (v) We selected 163,855 segments (95%) from the initial particle stack by one round of 2D classification with relion_refine (using 2-times binned data), discarding mostly segments that were extracted too close to the base or tip of the VSV bullet.

#### Initial 3D reconstruction

We obtained an initial 3D reconstruction from a subset of 871 segments selected from a visually good-looking 2D class. Using relion_reconstruct we prepared a first 3D reference by just applying the alignment parameter from the 2D alignment and C_1_ symmetry (using 4-times binned data). With auto-refinement in relion_refine (helical_inner_diameter = 260 Å, helical_outer_diameter = 700 Å, helical_nr_asu = 35, helical_z_percentage = 0.1, helical_twist_initial = -9.6°, helical_rise_initial = 1.379 Å) and imposing the previously reported helical symmetry (37.5 subunits per turn, helical pitch = 51.7 Å) (Ge et al., 2010), we obtained a 8.9 Å resolution reconstruction.

#### Helical reconstruction protocol

We set up an iterative helical reconstruction scheme using programs from Frealign (Grigorieff, 2016), cisTEM (Grant et al., 2018), RELION (Scheres, 2012), and EMAN2 (Bell et al., 2016) where each cycle involved the following steps: (i) References- based alignment of particle segments with refine3d (version 1.01) (first cycle in mode 3, subsequent cycles mode 1, C_1_ symmetry). (ii) 3D reconstruction of the full and half maps based in the alignment parameters with frealign_v9.exe (version 9.11) (C_1_ symmetry). Note that for reconstruction of the half maps, we used the first and second half of the particle segments (and not even and odd numbers, respectively) in order to prevent inflation of the half-map Fourier shell correlation (FSC) that would had resulted if segments extracted from the same trunk ended up contributing to different half maps. We also turned off Wiener filtering (FFILT = F) in order to avoid artifacts in the reconstructions, presumably because of the non-standard spectral signal-to-noise ratio of helical structures (e.g. layer planes in reciprocal space with very strong signal and no signal in between). (iii) Helical symmetry refinement with relion_helix_toolbox with the C_1_-symmetric map from the previous step as input (cyl_inner_diameter = 246–326 Å [depending on the number of subunits per turn], cyl_outer_diameter = 346–626 Å [depending on the number of subunits per turn], z_percentage = 0.3, sphere_percentage = 0.9). Symmetry search was local with a search range of 0.95–1.05 times the initial value for the helical rise, and of a helical twist corresponding to ±0.1 of the initial value of the number of subunits per turn. (iv) Real space helical symmetrization of the full and half maps with relion_helix_toolbox (z_percentage = 0.3, sphere_percentage = 0.9). (v) Masking of the helically-symmetrized maps with a cylindrical shell mask (inner radius = 123–163 Å, outer radius = 273–313 Å) with e2proc3d.py. (vi) FSC calculation from the half maps with e2proc3d.py. Applying this reconstruction protocol with the previously reported helical symmetry (Ge et al., 2010) to all 163,855 segment particles resulted in a map of about 7.5 Å resolution (using 2-times binned data).

#### Symmetry and geometry analysis

To determine the apparent rotational symmetry from top views of partially assembled or disrupted nucleocapsids (Fig. S1B, left), we calculated for each image the rotational self-correlation within a radius of 153–255 Å with “OR MAP” from SPIDER (Shaikh et al., 2008) (Fig. S1B, middle, blue curve). We numerically fit equation (1) to the calculated self-correlation *CC* within 5–60° of the self-rotation angle ϕ using SciPy (Virtanen et al., 2020) (Fig. S1B, right, red curve).

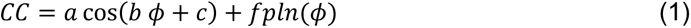

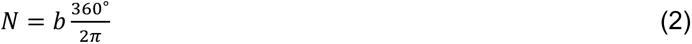

A third degree polynomial *fpln* was first obtained from an initial fit (Fig. S1B, middle, green curve), and the number of submits per turn *N*, or apparent rotational symmetry, is given by equation (2).

To measure the width and the length of VSV bullet particles, we extracted 2-times binned images (1400×1400 pixels) centered at the middle of each manually picked trunk. We used relion_autopick as a bilayer-enhancement algorithm by calculating a figure-of-merit image for each VSV bullet image based on a membrane reference image as input. For each particle that was fully within the extracted box, we manually measured its length, with e2helixboxer.py, by defining points centered on the membrane at the apex and bottom, respectively. To measure its width, we first projected density of the membrane-enhanced images along the particle axis, and then used Python to locate the two membrane peaks within the 1D-density distribution.

#### Supervised classification of helical segments

To prepare 3D references with different geometries for supervised classification of helical segments, we expanded the asymmetric unit density of the initial 7.5 Å-resolution reconstruction. We chose to describe flattening due to specimen preparation by modeling assemblies as elliptical helices. For this, we first rigid-body fitted into the 7.5 Å-resolution reconstruction density one asymmetric unit consisting of one N protein subunit with 9 RNA nucleotides from PDB-ID 2GIC (Green et al., 2006) and two M protein submits from PDB-ID 1LG7 (Gaudier et al., 2002), and then used the fitted models to generate a mask for density extraction. We defined a reference point within the extracted density, and a reference vector chosen normal to the tangential plane of the helix. Expansion then involved repetitive placement of the reference point for the asymmetric-unit density along the path of an elliptical helix while keeping the reference vector normal to the tangential plane of the elliptical helix (Figure 1B). The geometry of the resulting 3D references is defined by the geometry the elliptical helix and the number of subunits per helical turn. The pitch of the elliptical helix *P* is related to the helical rise *Δz* and the number of subunits per turn *N*:

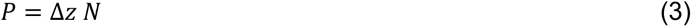

The major (long) and minor (short) axis of the elliptical helix, *a* and *b*, depend on the number of subunits per turn *N*, the distance along the helical path between reference points of adjacent asymmetric subunits (where *dxy* is its projection into the xy-plane), and the degree of degree of ellipticity (or flattening) *f*:

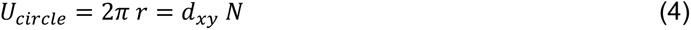

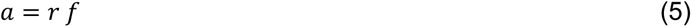

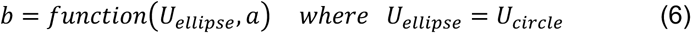

We used SciPy (Virtanen et al., 2020) for calculation of elliptical line integrals and to numerically determine *b* from the elliptical circumference *U_ellipse_* and *a*. We also used the Biopython module Bio.PDB for coordinate manipulations withing Python (Hamelryck and Manderick, 2003). In summary, the values of *P*, *N*, *d_xy_*, and *f* define each 3D reference.

We first used our subset of 871 segment particles and aligned them with refine3d (mode 3) to a set of non-flattened (*f* = 1.00) 3D references with constant pitch *P*, different numbers of subunits per turn (*N* = 34.5, 35.5, 36.5, 37.5, 38.5, 39.5, 40.5, 41.5), and a varying *d_xy_*. The pitch *P*, which we obtained by Fourier analysis of segments of trunk images, was kept constant, because it is determined by the axial stacking of helical rungs and does not depend on *N* or *f*. Class assignment (where each class corresponds to one particular 3D reference) was based on the highest alignment score and resembled a Gaussian distribution with respect to *N* (Figure 1C). For each *N*, distributions peaked sharply at a same value for *d_xy_*. A constant distance between adjacent subunits would be expected if the number of RNA nucleotides per N protein were constant among assemblies with different numbers of subunits per helical turn. We therefore kept the same value for *d_xy_* during subsequent analysis.

For classification of the full segment particle stack with 163,855 images, we relied on a high performance computer cluster that allowed us run parallel alignment calculations on up to 2,000 CPUs. Using 4-times binned data, we first globally aligned all images to eight non-flattened 3D references (*N* = 34.5, 35.5, 36.5, 37.5, 38.5, 39.5, 40.5, 41.5) using refine3d (mode 3). For each of the eight global alignments, we expanded the particle stack 39-fold according to the helical symmetry given by *N* using relion_particle_symmetry_expand. The expanded particle stack (6,390,345 images) was then locally aligned to 96 3D references using refine3d (mode 1), where we provided for each number of subunits per turn (*N* = 34.5, 35.5, 36.5, 37.5, 38.5, 39.5, 40.5, 41.5) twelve 3D references with different degrees of flattening (*f* = 1.000, 1.004, 1.007, 1.011, 1.015, 1.018, 1.022, 1.025, 1.029, 1.033, 1.036, 1.040). Class assignment and particle selection was again based on the highest alignment score with respect to the 3D references and expanded images, respectively (Figure 1C).

#### Helical reconstructions

We selected segment particles from the different *N*-classes without flattening and with only minimal fattening and applied for each particle stack a helical reconstruction protocol as outlined above using 2-times binned data. The number of helical segment particles ranged from 228 (*N* = 41.5) to 21,530 (*N* = 38.5) (Table S1). Helical reconstructions were then calculated with non-binned images and used for particle polishing with relion_motion_refine. After another round of the helical refinement protocol using the polished particle stacks, the resolution of the helical reconstructions, as determined by half-map FSC after masking, ranged from 12.1 Å (*N* = 41.5) to 4.1 Å (*N* = 38.5) (Table S1 and Figure S3A). Beam tilt correction did not further improve the resolution of the reconstructions. Local resolution was estimated with the program ResMap (Kucukelbir et al., 2014). We post-processed the *N* = 38.5 map by first sharpening it with cisTEM’s sharpen_map (Grant et al., 2018). The sharpened map was then used as input for LocScale (Jakobi et al., 2017), together with a preliminary refined model with estimated B factors. The resulting locally contrast-optimized map was again sharpened with sharpen_map.

#### Local reconstruction

To calculate a local reconstruction from the average of all asymmetric subunits (comprising one N protein, nine RNA bases, and two M proteins) captured in helical segments, we extracted subparticles as previously described (Herrmann et al., 2021; Jenni et al., 2019). For each *N* class, subparticles with a box size of 256×256 pixels were extracted around the central helical turn, because segment images were initially extracted with a one-turn offset. Subparticle stacks were signal-subtracted using relion_project and a mask that excluded the asymmetric subunit. The signal-subtracted stacks from the different *N*-classes were aligned with respect to each other based on transformation matrices determined from rigid-body fitted PDB models. The resulting subparticle stack, containing approximately three million images, was further aligned by classification without alignment using three iterations of the following protocol: classification with cisTEM’s refine3d (8 classes, 5 Å high-resolution limit for classification, 60 cycles, 3D mask covering the asymmetric unit), superposition of the maps corresponding to the 8 classes based on rigid-body fitted models, and updating of the particle alignment parameters according to the 3D map alignment. As a final step, we carried out local alignment with refine3d (5 Å high-resolution limit for refinement, 8 cycles, 3D mask covering the asymmetric unit). The masked local reconstruction of the asymmetric unit, calculated from a non-signal-subtracted particle stack, had a nominal resolution of 3.5 Å (Table S1 and Figure S3B). We post-processed the map by sharpening it with sharpen_map (Grant et al., 2018) and local filtering with LocScale (Jakobi et al., 2017).

#### VSV tip reconstruction

To obtain a reconstruction of the VSV bullet tip, we initially selected 5,763 projection views of viral particles from the micrographs, in which we manually marked the base and tip of each bullet (Figure S6A). A segment extracted at the center of each particle was used to determine *N*, the number of subunits per turn in the helical region of the stalk, by supervised classification as described above (without talking flattening into account), yielding 2,521 particles with *N* of 38.5 (Figure S6B). The alignment determined at the center of the particles was then “shifted” towards the tips, to the point before the helical part transitions into narrowing turns. We did this stepwise according to helical symmetry with increments of ten, five and finally one asymmetric unit shifts, until the distance between the projection of a reference point on the z axis within the box of the 3D reference and the projection of the manually marked tip on the projection of the z axis in each particle image was minimal (Figure S6C and Movie S1). At each shift, we updated the alignment by local refinement with cisTEM’s refine3d. We then further aligned the tip images using classification without alignment (3 cycles 2D classification, and 1 cycle 3D classification). For this, we calculated translational offsets along the z axis between the individual classes from rotational averages and applied corresponding shifts according to helical symmetry (Figure S6D and E). Next, we symmetry expanded each tip image (+/-80 asymmetric subunits along the helical symmetry) and calculated scores using a 3D reference, where we rotationally averaged the membrane region, and which was reconstructed without the particle image for which the score was calculated. (Figure S6F, helical offset alignment cycle). This helical offset alignment was iterated for about 9 cycles until convergence (Figure S6G). As the reconstruction improved, we rigid-body fitted the structures of individual modules (N and M1 subunits) where the density was good enough to warrant an unambiguous fit. The fitted modules where then used to calculate a mask and transformation matrices for local symmetry density averaging. The resulting reconstruction was then used for particle picking in the original micrographs using cisTEM’s match_template, refine_template, and make_template_result (Lucas et al., 2021) (Figure S6H), yielding a stack of 25,401 images, which were then aligned with cisTEM (mode 3, global alignment, C1 symmetry) (Figure S6F, big cycle with 2D template matching).

### Model Building and Refinement

Templates for modeling the N protein were taken from PDB-IDs 2GIC (Green et al., 2006) and 5UK4 (Hanke et al., 2017). The M proteins where modeled based on PDB-ID 1LG7 (Gaudier et al., 2002). We used the programs O (Jones et al., 1991) for model building and adjustments, phenix.real_space_refine (Afonine et al., 2018) for structure refinement (standard stereochemical and *B*-factor restraints, as well as Ramachandran, rotamer, and secondary structure restraints), phenix.mtriage for FSC calculations between maps and models (Afonine, 2017), and MolProbity (Chen et al., 2010) to validate the models (Table S1).

A model that was refined against the post-processed local reconstruction map had the following composition, where the protomer indices refer to the numbering in the *N* = 38.5 helix: 15 RNA nucleotides modeled as poly-uridine (2 from N_i-1_, 9 from N_i_, and 2 from N_i+1_); nucleoprotein (N) residues 2–35 (N-terminal arm from N_i+1_), 20–341 and 366–422 (N- and C-terminal lobes from N_i_), 335–373 (extended loop from N_i-1_); matrix protein (M1) residues 43–59 (self-association motif from M1_i+38_), 57–227 (globular domain from M1_i_); matrix protein (M2) residues 58–229 (globular domain from M2_i_). Compared to the N protein structures of PDB-IDs 2GIC and 5UK4, we corrected a sequence register shift at residues 154–181. The FSC between the refined model and final local reconstruction map was 0.5 at a spatial frequency corresponding to a resolution of 3.9 Å (Figure S3B), suggesting that the 3.5 Å nominal resolution determined from the half maps was slightly overestimated.

We also fit the model from the local reconstruction into the 4.1 A-resolution map of the *N* = 38.5 reconstruction. For refinement with phenix.real_space_refine, we placed 15 copies of the asymmetric unit by stacking three partial turns with five consecutive units each. This allowed the central asymmetric unit to form all potential inter-subunit contacts during refinement. The asymmetric unit of the refined helical structure had the following composition: 9 RNA nucleotides modeled as poly-uridine; nucleoprotein (N) residues 2–422; matrix protein (M1) residues 43–227; matrix protein (M2) residues 58–229. The FSC between the refined model and final helical reconstruction map with *N* = 38.5 was 0.5 at a spatial frequency corresponding to a resolution of 4.6 Å (Figure S3C).

The refined structure of the asymmetric unit from *N* = 38.5 was then fit into the maps of the other different *N* classes by rigid-body fitting (Figure 2A). Each subunit was treated as a rigid-body, except in case of the low-resolution map from *N* = 41.5, were the entire asymmetric unit was defined as rigid body.

### Figure Preparation

We prepared the figures using PyMol (The PyMOL Molecular Graphics System, Version 2.1 Schrödinger, LLC) and matplotlib (Hunter, 2007). Amino acid sequences (Data S1–3) were aligned with MAFFT (Katoh et al., 2002) and multiple sequence alignment were displayed and annotated with ESPript (Robert and Gouet, 2014). Secondary structure assignments were calculated with DSSP (Kabsch and Sander, 1983). We used the PISA program to analyze subunit interfaces (Krissinel and Henrick, 2007). We obtained positional conservation scores from the multiple sequence alignments with the program ConSurf (Ashkenazy et al., 2016).

### Density Maps and Atomic Coordinates Accession Identifiers

The cryo-EM maps can be obtained from the Electron Microscopy Data Bank with accession identifiers EMD-26603 (local reconstruction) and EMD-26602 (*N* = 38.5 helical reconstruction). The refined atomic coordinates can be obtained from the Protein Data Bank with accession identifiers PDB-ID 7UML (local reconstruction) and PDB-ID 7UMK (*N* = 38.5 helical reconstruction).

## ACKNOWLEDGMENTS

We thank Z. Li, S. Sterling, R. Walsh and S. Rawson for facilitating our use of the Harvard Medical School Cryo-EM Center for Structural Biology and the Harvard Medical School Molecular Electron Microscopy Suite; the Nancy Lurie Marks Family Foundation for support of the Cryo-EM Center; the SBGrid Advanced Research Computing group for IT support, and T. Kirchhausen for access to computational resources. Portions of this research were conducted on the O2 High Performance Compute Cluster, operated by the Research Computing Group at Harvard Medical School. We acknowledge support from NIH grants R37-AI1059371 (to SPJW) and R01-CA13202 (to SCH). SCH is an Investigator in the Howard Hughes Medical Institute.

---

**Table S1.** Cryo-EM Data Collection and Model Statistics.

| Vesicular stomatitis virus (VSV, Indiana serotype) |  |  |  |  |  |  |  |  |
| --- | --- | --- | --- | --- | --- | --- | --- | --- |
| **Data collection** |  |  |  |  |  |  |  |  |
| Electron microscope | Titan Krios |  |  |  |  |  |  |  |
| Magnification | 58823 |  |  |  |  |  |  |  |
| Voltage (kV) | 300 |  |  |  |  |  |  |  |
| Defocus range ($\mu\text{m}$)<sup>a</sup> | 1.0–3.0 | | | | | | | |
| Pixel size ($\text{\AA}$) | 0.85 | | | | | | | |
| Number of movies | 18353 |  |  |  |  |  |  |  |
| **Helical reconstruction** | **N =** | **N =** | **N =** | **N =** | **N =** | **N =** | **N =** | **N =** |
|  | **34.5** | **35.5** | **36.5** | **37.5** | **38.5** | **39.5** | **40.5** | **41.5** |
| Number of images | 1274 | 5393 | 17480 | 20367 | 21530 | 10369 | 1509 | 228 |
| Box size (pixels) | 1200 | 1200 | 1200 | 1200 | 1200 | 1200 | 1200 | 1200 |
| Symmetry imposed | Helical | Helical | Helical | Helical | Helical | Helical | Helical | Helical |
| Helical twist ($^{\circ}$) | -10.44 | -10.14 | -9.86 | -9.60 | -9.35 | -9.11 | -8.89 | -8.67 |
| Helical rise ($\text{\AA}$) | 1.488 | 1.449 | 1.411 | 1.374 | 1.341 | 1.309 | 1.276 | 1.248 |
| Refinement mask diameter ($\text{\AA}$)<sup>b</sup> | 246, 546 | 258, 558 | 268, 568 | 280, 580 | 292, 592 | 302, 602 | 314, 614 | 326, 626 |
| FSC mask diameter ($\text{\AA}$)<sup>b</sup> | 274, 534 | 286, 546 | 296, 556 | 308, 568 | 320, 580 | 330, 590 | 342, 602 | 354, 614 |
| Map resolution ($\text{\AA}$)<sup>c</sup> | 6.9 | 4.9 | 4.3 | 4.2 | 4.1 | 4.5 | 6.6 | 12.1 |
| **Local reconstruction** |  |  |  |  |  |  |  |  |
| Number of images | 2985175 |  |  |  |  |  |  |  |
| Box size (pixels) | 256 |  |  |  |  |  |  |  |
| Symmetry imposed | C<sub>1</sub> |  |  |  |  |  |  |  |
| Map resolution ($\text{\AA}$)<sup>c</sup> | 3.5 | | | | | | | |
| **Model statistics** | **Local reconstruction** |  |  |  | **Helical reconstruction N = 38.5** |  |  |  |
| EMD accession identifier | EMD-26603 |  |  |  | EMD-26602 |  |  |  |
| PDB accession identifier | 7UML |  |  |  | 7UMK |  |  |  |
| Refinement resolution ($\text{\AA}$) | 3.8 | | | | 4.1 | | | |
| CC (mask) | 0.70 |  |  |  | 0.75 |  |  |  |
| Model composition |  |  |  |  |  |  |  |  |
| Non-hydrogen atoms | 6750 |  |  |  | 6408 |  |  |  |
| Protein residues | 812 |  |  |  | 778 |  |  |  |
| RNA nucleotides | 13 |  |  |  | 9 |  |  |  |
| B factors ($\text{\AA}^2$) | | | | | | | | |
| Protein | 44.6 |  |  |  | 157 |  |  |  |
| RNA | 33.5 |  |  |  | 138 |  |  |  |
| R.m.s. deviations |  |  |  |  |  |  |  |  |
| Bond lengths ($\text{\AA}$) | 0.003 | | | | 0.002 | | | |
| Bond angles ($^{\circ}$) | 0.575 | | | | 0.534 | | | |
| Validation |  |  |  |  |  |  |  |  |
| MolProbity clash score | 7.4 |  |  |  | 5.2 |  |  |  |
| Poor rotamers (%) | 0.0 |  |  |  | 0.0 |  |  |  |
| Ramachandran plot |  |  |  |  |  |  |  |  |
| Favored (%) | 97.4 |  |  |  | 97.9 |  |  |  |
| Allowed (%) | 2.0 |  |  |  | 1.6 |  |  |  |
| Disallowed (%) | 0.6 |  |  |  | 0.5 |  |  |  |

<sup>a</sup> Approximate range of underfocus.<sup>b</sup> Inner and outer diameter of the cylindrical masks used for refinement and FSC calculation, respectively.<sup>c</sup> Resolution where FSC between masked half maps drops below 0.143.

**Figure S1.**
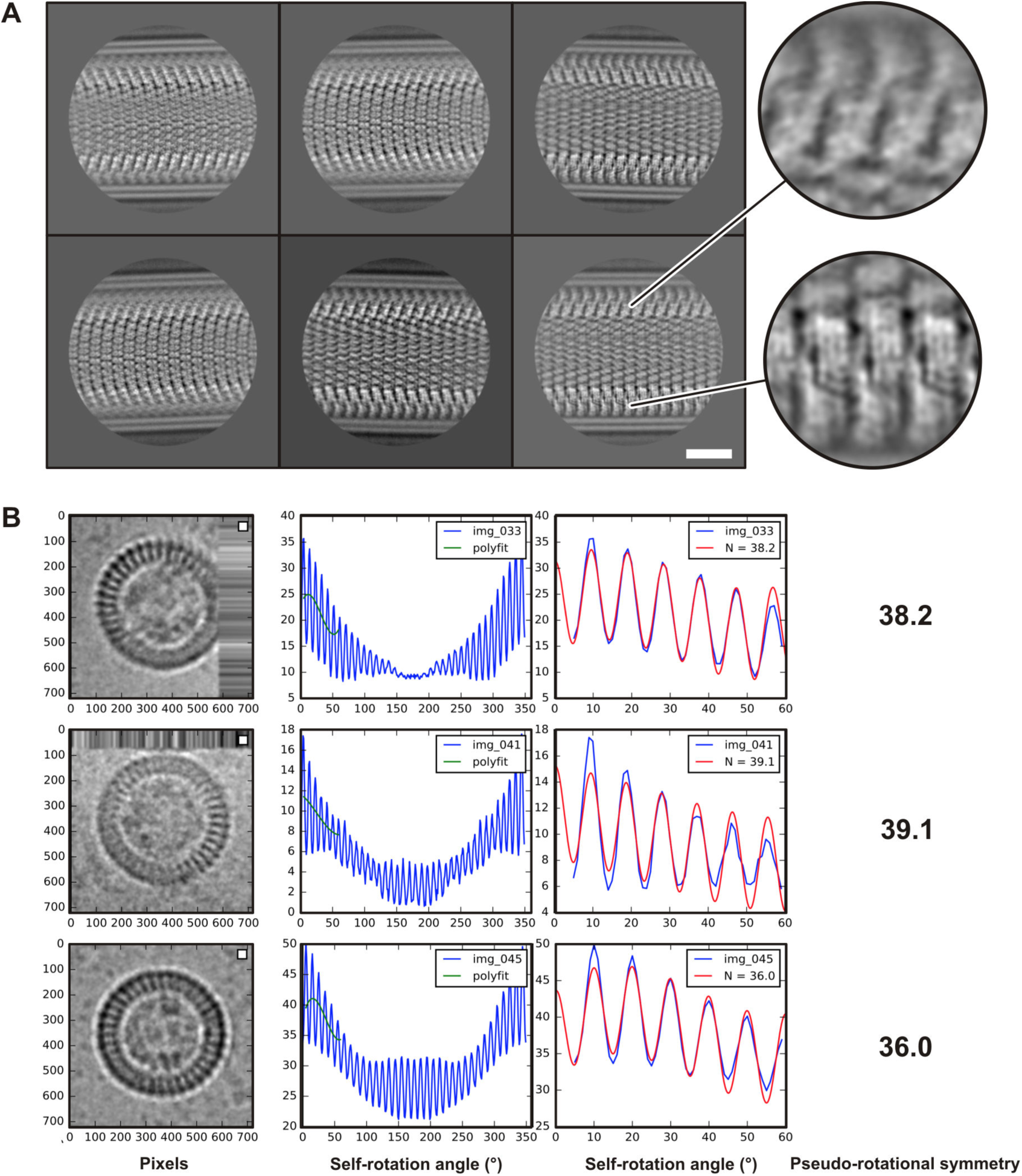
Heterogeneity of VSV Helical Nucleocapsid Segments. (A) Representatives of best looking 2D class averages. Magnifications of two different regions of the bottom right class average show different degrees of sharpness, indicating structural heterogeneity and uneven focusing of the 2D alignment. The scale bar corresponds to 20 nm. (B) Three top-view images of partially assembled or disrupted virion fragments. We determined the apparent numbers of subunits per turn by fitting a cosine function (red curve) to the rotational self-correlation (blue curve) obtained from the observed density distribution within a radius of 153–255 Å. The green curve is a polynomial fit used for base line estimation.

**Figure S2.**
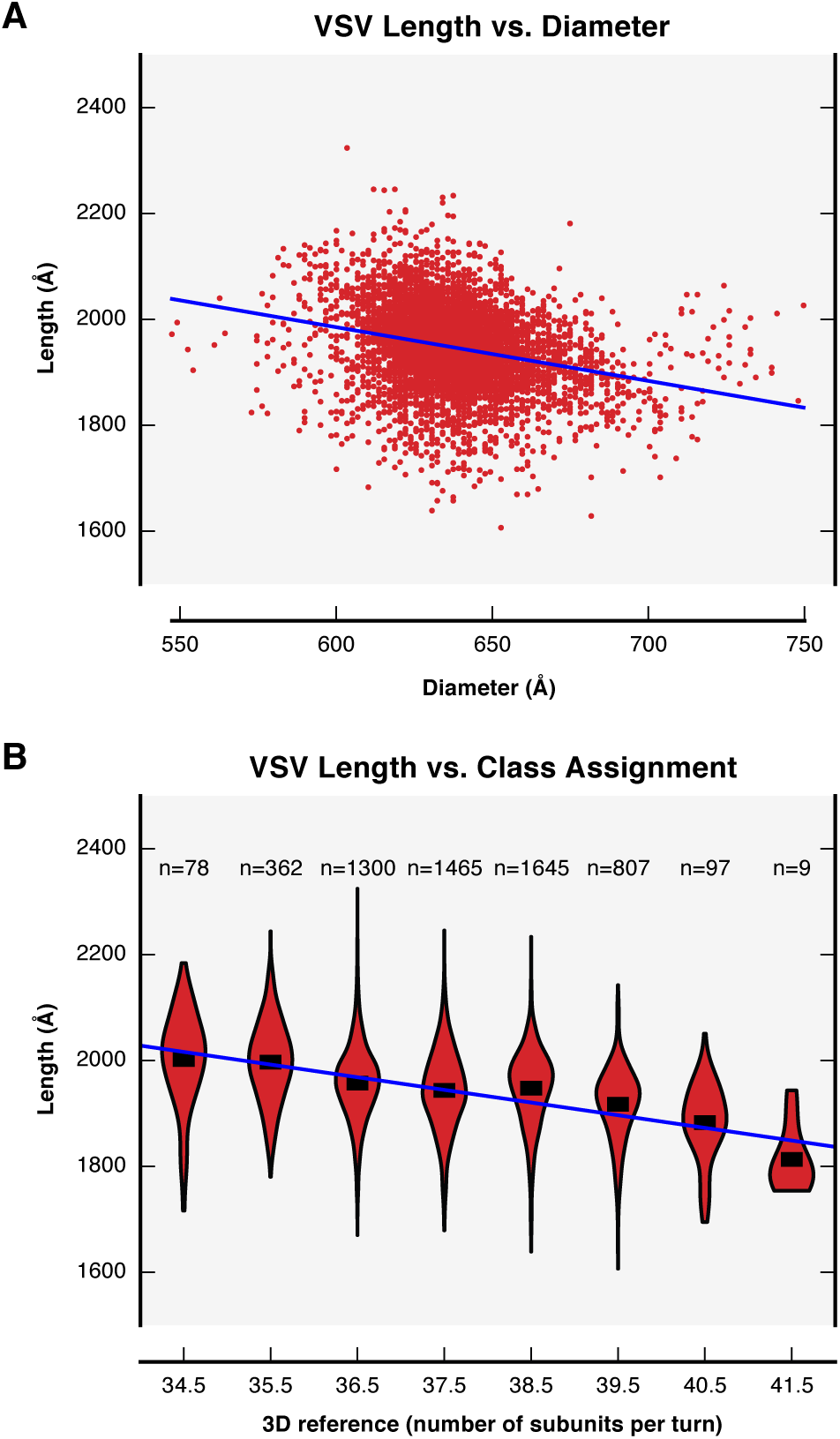
Structural Heterogeneity of VSV Virions. (A) Scatter plot of measured length versus diameter of individual VSV virions. The blue line is a linear regression fit to the data points (R-squared = 0.07, slope = -1.0). (B) Violin plot of measured length versus class assignment (after supervised classification) of individual virions. Length distributions and the number of virions and are shown for each class. Means are shown as horizontal black bars. The blue line is a linear regression fit to the means (R-squared = 0.89, slope = -23.9 per class).

**Figure S3.**
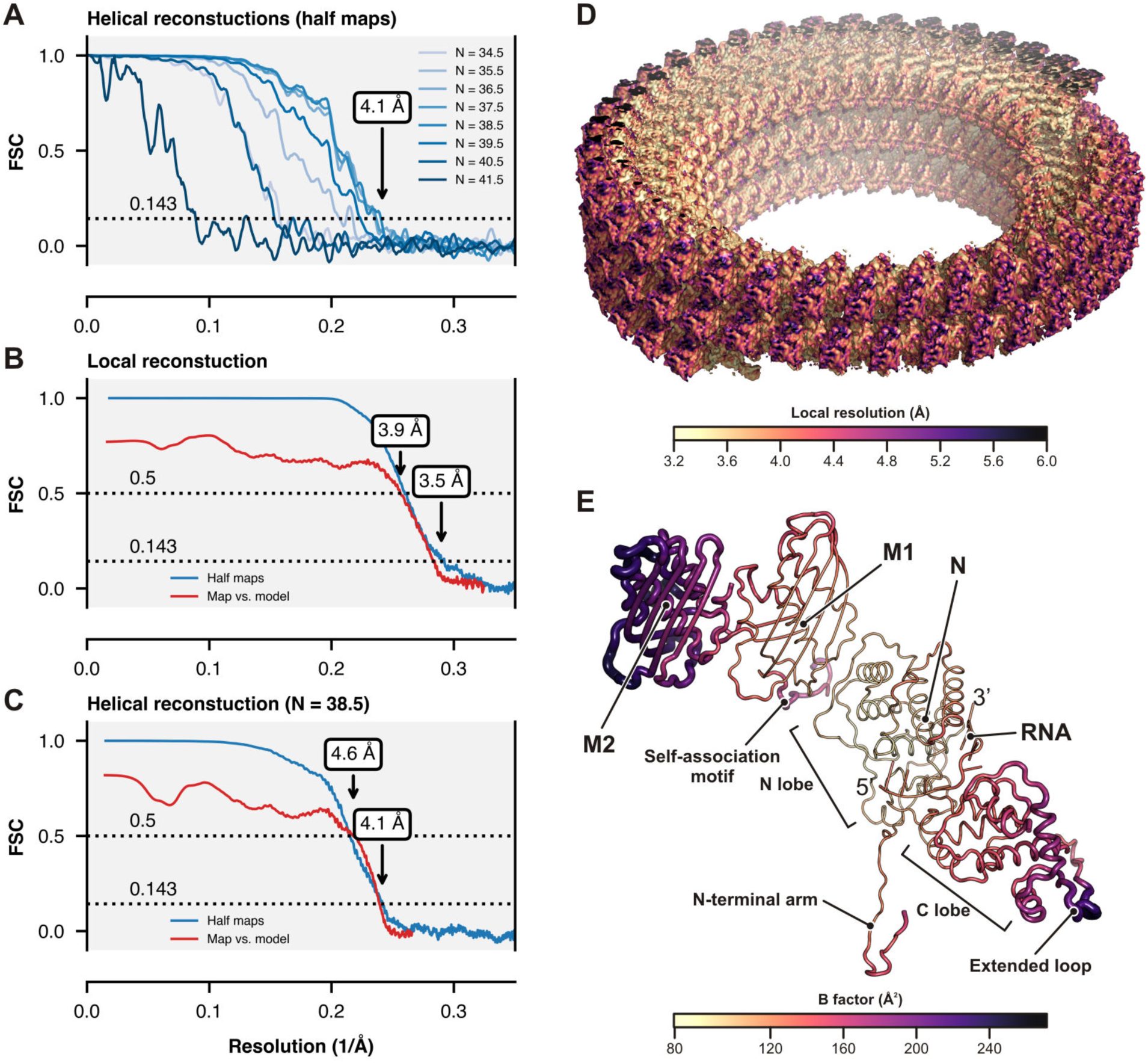
Resolution Estimation of Cryo-EM Reconstructions. (A) Top, Fourier shell correlation (FSC) between the two half maps for each of the helical reconstructions with different numbers of subunits per turn (*N*). Symmetrized half maps were masked with hollow cylindrical masks with inner and outer diameter as shown in Table S1. The overall resolution where the correlation drops below 0.143 is 4.1 Å for the reconstruction with *N* = 38.5. (B) FSC analysis of the local reconstruction. The blue curve is the FSC between the half maps after masking based on the refined model. The red curve is the FSC between the final map and the refined model. The overall resolution is 3.5 Å for the half maps where the correlation drops below 0.143 and 3.8 Å for the model where the correlation drops below 0.5. (C) FSC analysis of the *N* = 38.5 helical reconstruction. The blue curve is the FSC between the half maps after masking based on the refined model. The red curve is the FSC between the final map and the refined model. The overall resolution is 4.1 Å for the half maps where the correlation drops below 0.143 and 4.6 Å for the model where the correlation drops below 0.5. (D) Local resolution estimation color-mapped on the *N* = 38.5 helical reconstruction. (E) *B* factors mapped on the refined structure of the *N* = 38.5 helical reconstruction. Note that that *B* factor values are only meaningful relative within the structure as they depend on the degree of sharpening that was applied to the cryo-EM reconstruction.

**Figure S4.**
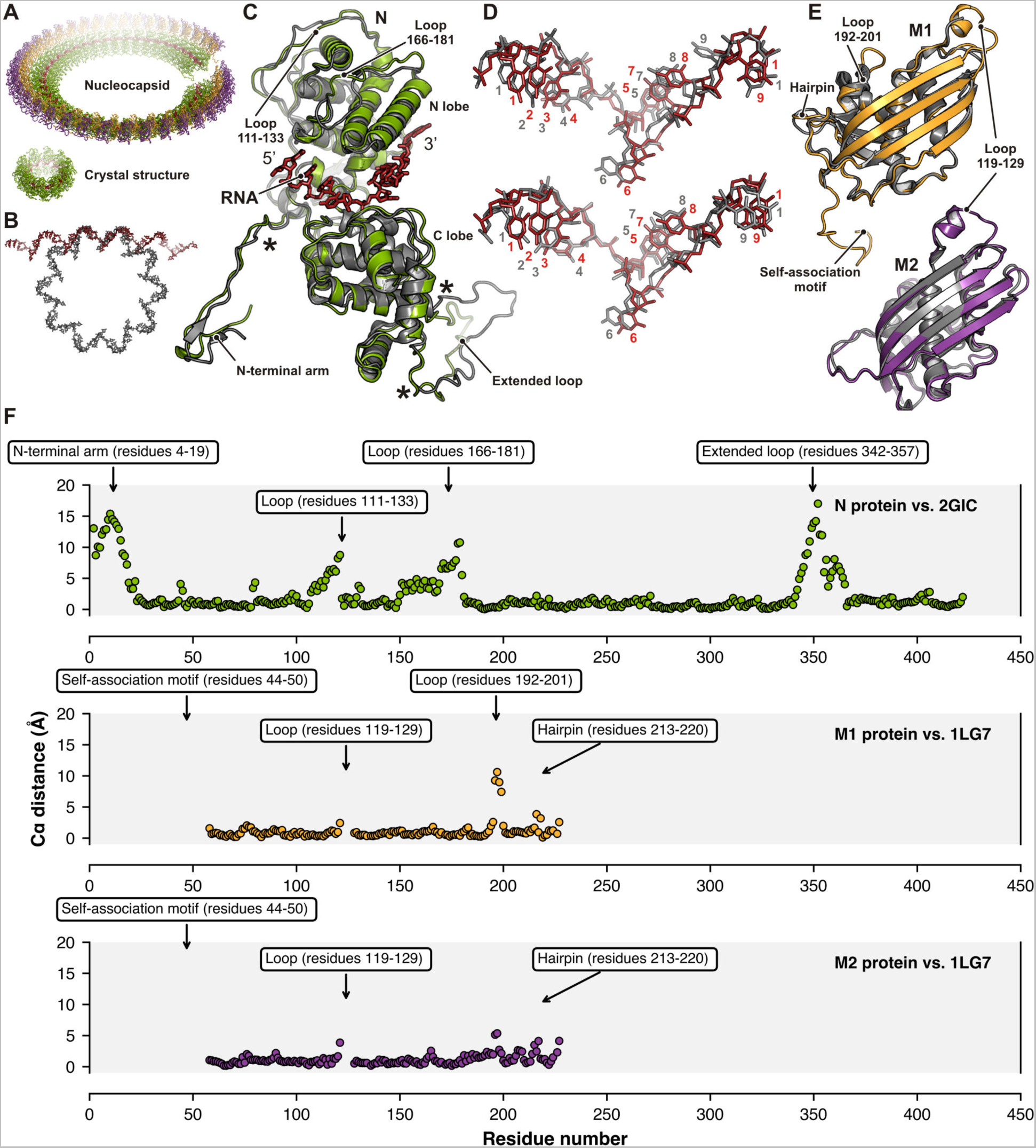
Structural Comparison of Nucleocapsid-Assembled and Crystallized RNA, N, and M Proteins. (A) Overviews of the nucleocapsid (top, one turn of the N = 38.5 structure is shown) and crystallized decameric RNP complex (bottom) are shown. RNA is colored in red, N protein in green, M1 and M2 proteins in orange and purple, respectively. (B) Comparison of the RNA as observed in the nucleocapsid (red) and the crystal structure (gray). (C) Superposition of the N protein from the nucleocapsid (green) and from the crystal structure (gray, PDB-ID 2GIC). Segments with substantially different conformations are labelled. Asterisks indicated flexibility points that allow the N-terminal arm and the extended loop to adjust for the different packing in the nucleocapsid and crystal structure. Residues of loop 111– 133, solvent exposed or in lattice contacts in the crystal structures, interact with the M1 layer in the assembled nucleocapsid. Loop 166–181, mostly solvent exposed in the nucleocapsid, was incorrectly modeled in the crystal structures PDB-IDs 2GIC and 5UK4 (a sequence register shift). (D) Close-up view of the RNA structure from one repeating unit from the nucleocapsid (red) and the crystal structures (gray) PDB-ID 2GIC (top) and on PDB-ID 5UK4 (bottom) after superposition of the N proteins. Nucleotides are labeled 1–9. Note the different conformation of nucleotide 9. (E) Superposition of the M1 and M2 proteins from the nucleocapsid (orange and purple) and from the crystal structure (gray, PDB-ID 1LG7). Segments with substantially different conformations are labelled. The poorly ordered loop 119–129 in our density maps was not modeled in the crystal structure. M1 loop 192–201 and hairpin 213–220 bind the C terminus of the M2 subunits in the nucleocapsid. (F) The plots show the distances between corresponding Cα atoms after superposition of the N protein (green, calculated from residues 27–341 and 372–422) on PDB-ID 2GIC; the M1 protein (orange, calculated from residues 58–121 and 128-227) on PDB-ID 1LG7; the M2 protein (purple, calculated from residues 58–121 and 128-227) PDB-ID 1LG7. Regions with large conformational shifts are labeled on the top of each plot.

**Figure S5.**
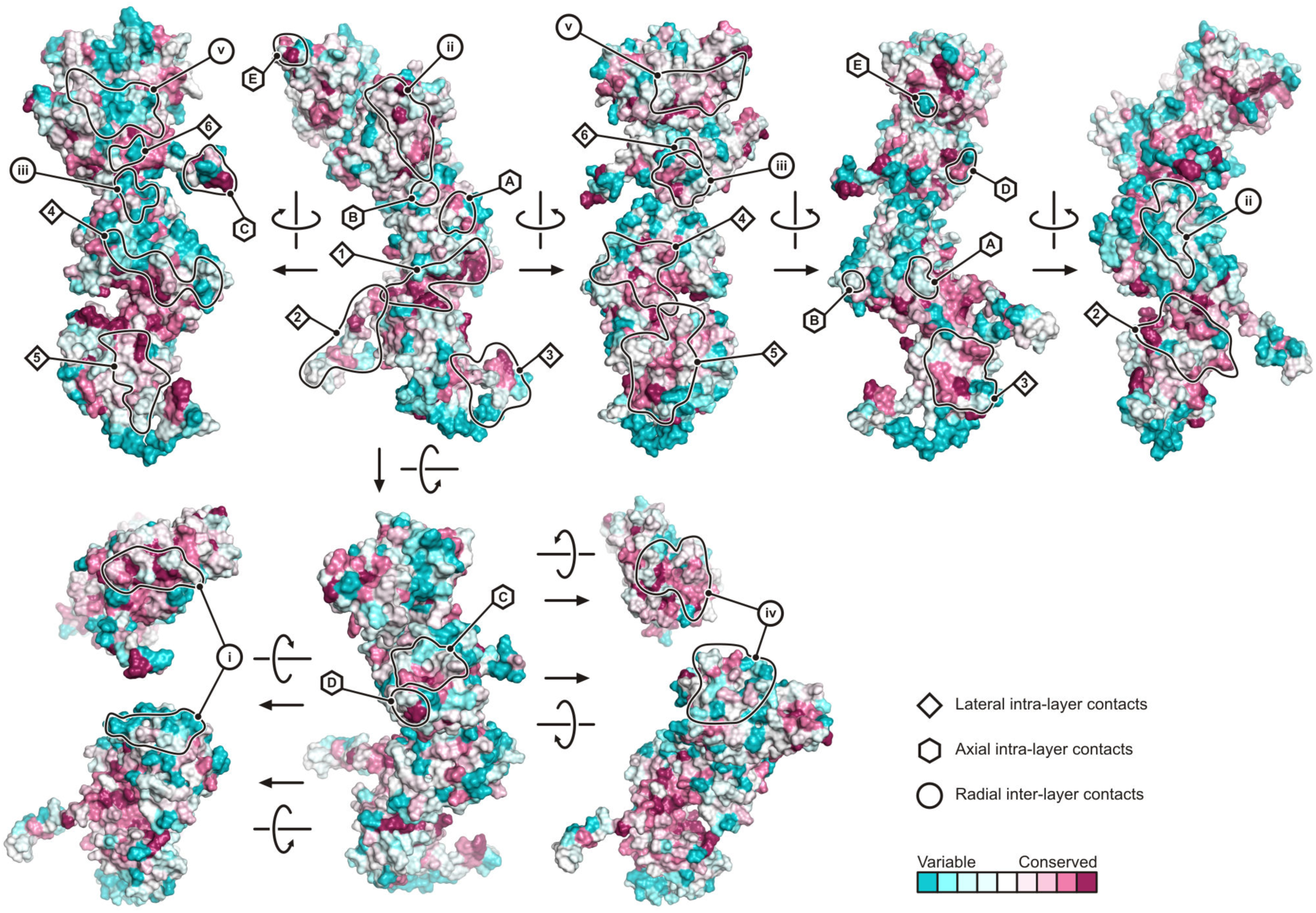
Conservation Analysis of Inter-Subunit Contacts. The N, M1, and M2 subunits of one module are shown in surface representation and colored according to amino acid conservation (see Data S1–S3 for multiple sequence alignments). Patches that form inter-subunit contacts are indicated and labeled as in Figure 4.

**Figure S6.**
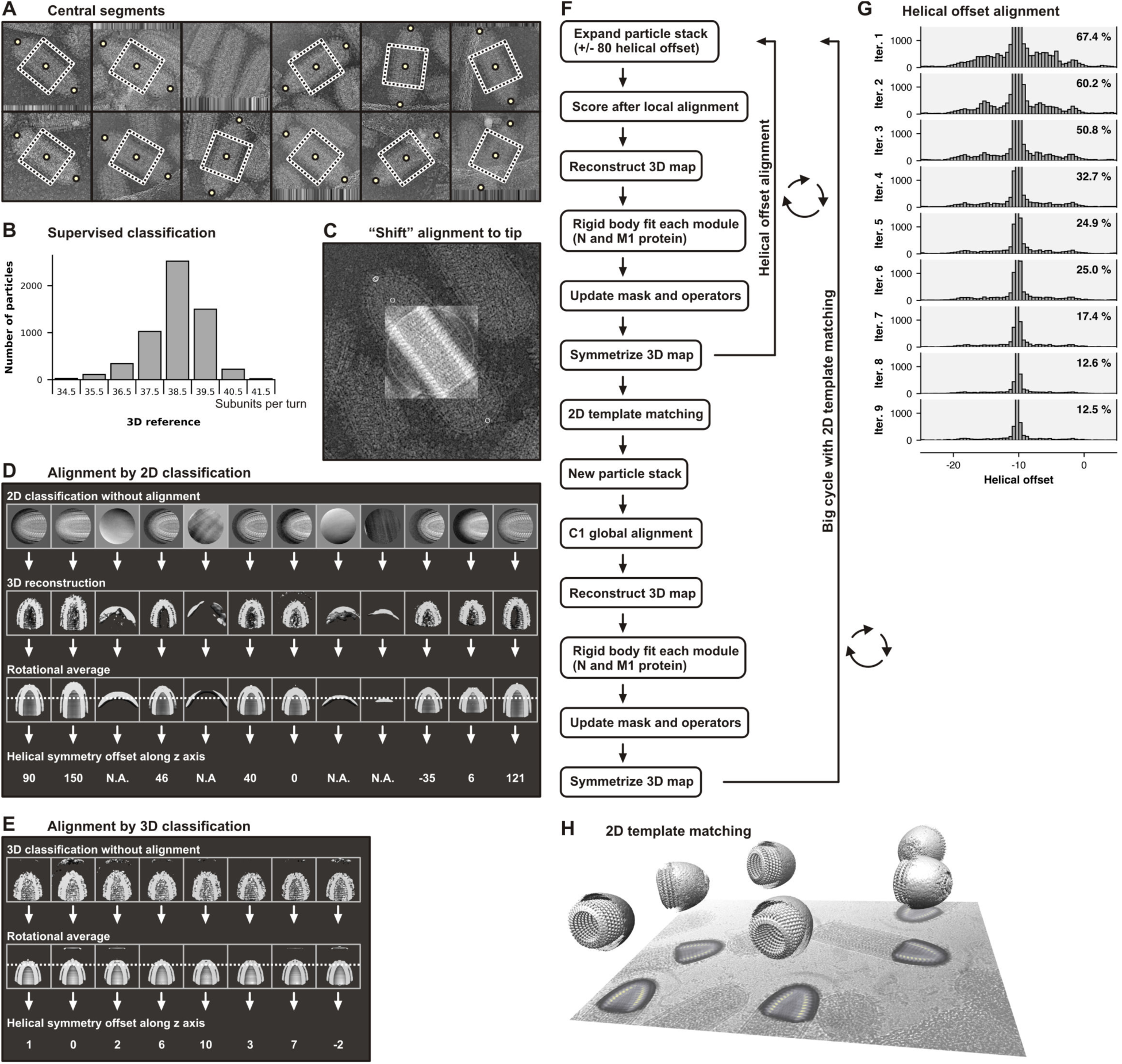
Reconstruction of the VSV Tip. (A) Selection of central segment for initial alignment. The position for segment extraction was obtained after manually marking the top and bottom of virions in the micrographs (yellow dots). (B) Supervised classification of central segments. Only non-flattened 3D references with different numbers of subunits per turn were used for the analysis. (C) Shift of the alignment towards the VSV tip (see Movie S1). (D) Alignment by 2D classification (without alignment). (E) Alignment by 3D classification (without alignment). (F) Flow chart showing the individual steps used to refine the reconstruction of the tip. The large cycle, which includes 2D template matching and global alignment, is computationally expensive. The small cycle (helical offset alignment) is much faster than the large cycle and was iterated to convergence. (G) Convergence of the small cycle (helical offset alignment). The histograms show the distribution of particles with the best match to the current reference after applying a corresponding helical offset for each iteration. The percentage of particles with a helical offset is shown for each iteration. (H) 2D template matching illustrated for a representative micrograph.

**Figure S7.**
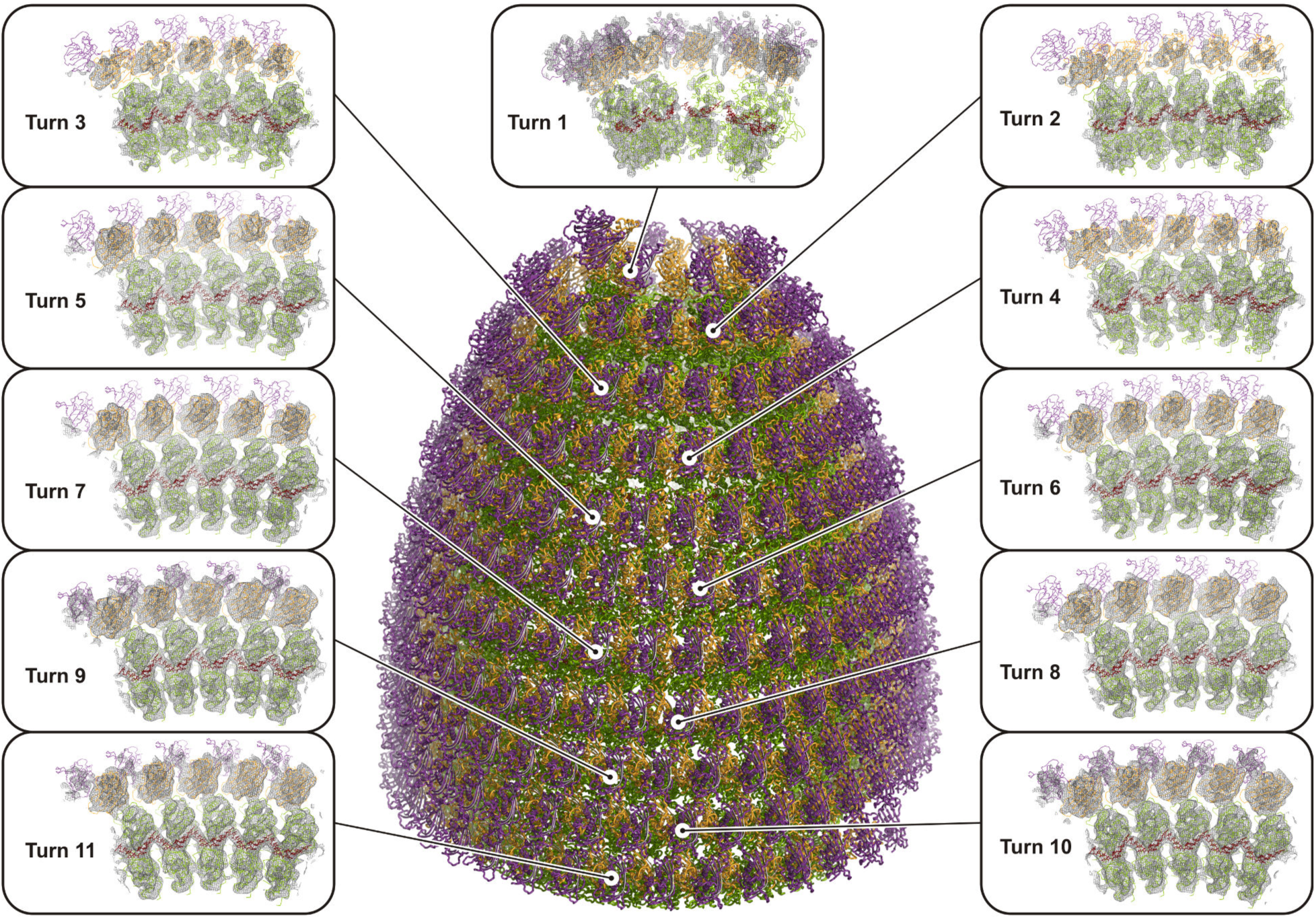
Structure of the VSV Tip. In the center, a side view of the VSV tip structure is shown in ribbon representation. The nucleoprotein (N) is colored green, the RNA red, and the two matrix proteins (M1 and M2) are colored orange and purple, respectively. Modules comprised of one N, M1, and M2 proteins, and 9 nucleotides were fitted as rigid bodies. On the left and right, fits of the modules in the tip reconstruction density are shown for the first 11 turns of the RNP ribbon.

**Figure S8.**
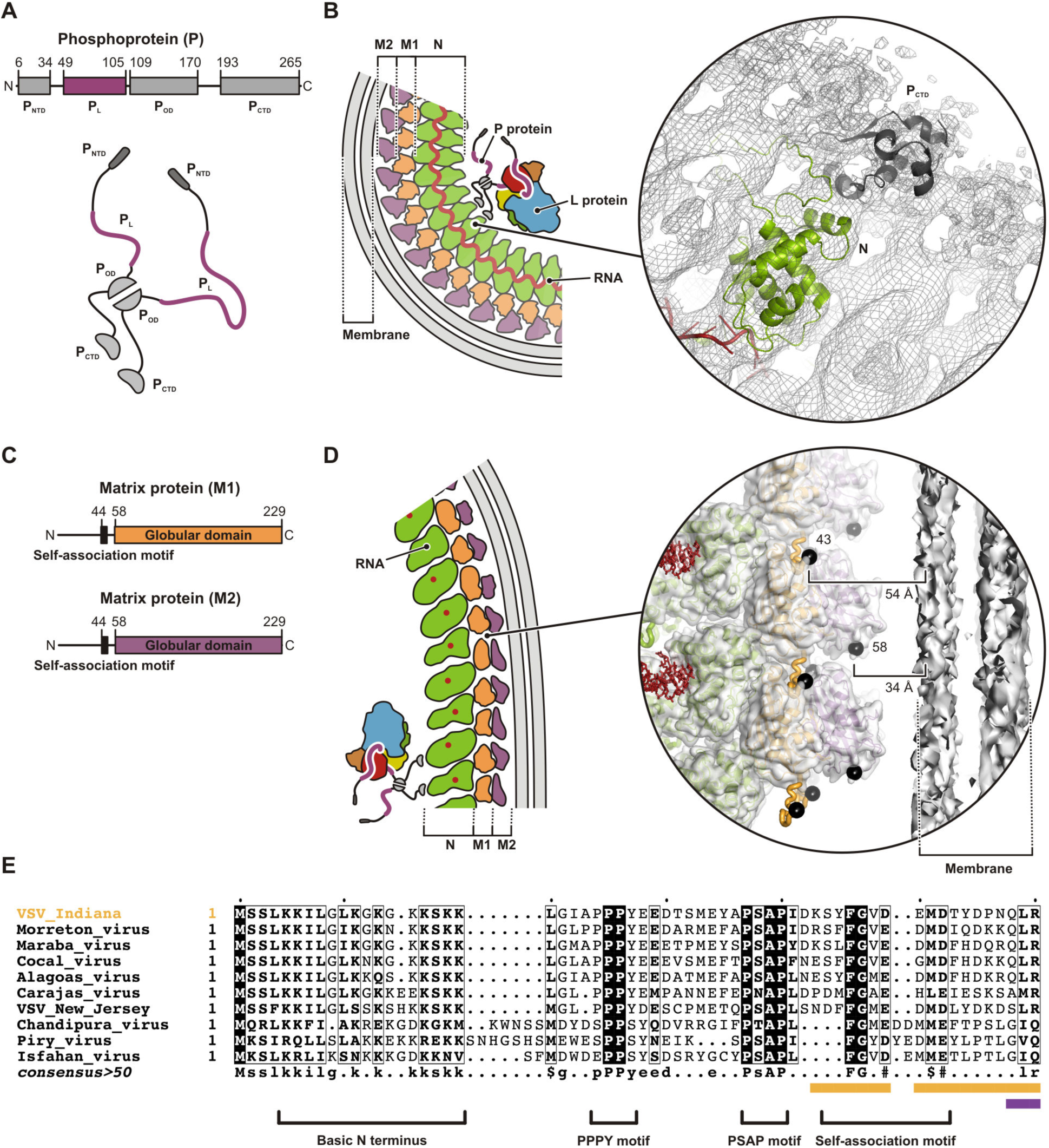
Binding of the Phosphoprotein to the RNP and Potential Membrane-Binding of the M protein. (A) Top, linear domain organization of the phosphoprotein (P). Amino acid numbers indicate domain boundaries. The P protein’s N-terminal (P_NTD_), oligomerization (P_OD_), and C-terminal (P_CTD_) domains are in gray; its L protein-binding domain (P_L_) is in magenta. Bottom, schematic drawing of a P-protein dimer. (B) Density of the P protein’s C-terminal (P_CTD_, gray) domain bound to the N protein (green) was observed in low-pass filtered maps at low contour levels. (C) Linear domain organization of the M1 and M2 proteins (as in Figure 3A). (D) Distance between the first modelled residues of the M proteins (43 for M1, and 58 for M2) and the membrane of the VSV helical trunk. (E) Multiple sequence alignment of the M protein N terminus (residues 1–160 as in Data S1). Modeled residues of the M1 and M2 are indicated at the bottom by orange and purple bars, respectively. Sequence motifs of interest are labeled.

## Data S1. N Protein Multiple Sequence Alignment

N protein amino acid sequences from different viruses were obtained from UniProt with their sequence accession identifiers in parenthesis: VSV_Indiana (P03523), vesicular stomatitis virus (Indiana strain); Morreton_virus (A0A0D3R1D1), Morreton vesiculovirus; Maraba_virus (F8SPF5), Maraba virus; Cocal_virus (B3FRL0), Cocal virus; Alagoas_virus (B3FRL5), vesicular stomatitis Alagoas virus; Carajas_virus (A0A0D3R1M8), Carajas virus; VSV_New_Jersey (P16379), vesicular stomatitis New Jersey virus; Chandipura_virus (P11211), Chandipura virus; Piry_virus (A0A1I9L1X2), Piry virus; Isfahan_virus (P16379), Isfahan virus; Rabies_virus (P06025), Rabies virus; Mokola_Virus (P0C570), Mokola virus. Assigned secondary structure elements are annotated on top of the sequences. N protein residues that are within a distance of 5 Å to the RNA or neighboring N and M1 subunits in the structure of the *N* = 3.85 helical reconstruction are labeled underneath the sequences. Residues that contact the RNA are labeled with red ovals. Residues that contact N and M1 proteins are labeled with green and orange ovals, respectively.

## Data S2. M1 Protein Multiple Sequence Alignment

M1 protein amino acid sequences from different viruses were obtained from UniProt with their sequence accession identifiers in parenthesis: VSV_Indiana (P03523), vesicular stomatitis virus (Indiana strain); Morreton_virus (A0A0D3R1D1), Morreton vesiculovirus; Maraba_virus (F8SPF5), Maraba virus; Cocal_virus (B3FRL0), Cocal virus; Alagoas_virus (B3FRL5), vesicular stomatitis Alagoas virus; Carajas_virus (A0A0D3R1M8), Carajas virus; VSV_New_Jersey (P16379), vesicular stomatitis New Jersey virus; Chandipura_virus (P11211), Chandipura virus; Piry_virus (A0A1I9L1X2), Piry virus; Isfahan_virus (P16379), Isfahan virus. M1 protein residues that are within a distance of 5 Å to neighboring N, M1, and M2 subunits in the structure of the *N* = 3.85 helical reconstruction are labeled underneath the sequences. Residues that contact N proteins are labeled with green ovals. Residues that contact M1 and M2 proteins are labeled with orange and purple ovals, respectively.

## Data S3. M2 Protein Multiple Sequence Alignment

M2 protein amino acid sequences from different viruses were obtained from UniProt with their sequence accession identifiers in parenthesis: VSV_Indiana (P03523), vesicular stomatitis virus (Indiana strain); Morreton_virus (A0A0D3R1D1), Morreton vesiculovirus; Maraba_virus (F8SPF5), Maraba virus; Cocal_virus (B3FRL0), Cocal virus; Alagoas_virus (B3FRL5), vesicular stomatitis Alagoas virus; Carajas_virus (A0A0D3R1M8), Carajas virus; VSV_New_Jersey (P16379), vesicular stomatitis New Jersey virus; Chandipura_virus (P11211), Chandipura virus; Piry_virus (A0A1I9L1X2), Piry virus; Isfahan_virus (P16379), Isfahan virus. M2 protein residues that are within a distance of 5 Å to neighboring M1 and M2 subunits in the structure of the *N* = 3.85 helical reconstruction are labeled underneath the sequences. Residues that contact M1 and M2 proteins are labeled with orange and purple ovals, respectively.

## Movie S1. VSV Tip Reconstruction

Shift of the alignment towards the VSV tip. A projection of the 3D reference according to the alignment parameters is mapped onto the virion image. The two small circles at the virion tip are (i) the tip initially marked manually in the micrographs and (ii) its projection onto the projection of the helical axis of the aligned 3D reference. The small circle with a dot is the projected center of the reference box. The small circle in between is a projected defined point on the z axis (helical axis) of the reference box. The movie shows how the alignment is shifted from an initial central segment towards the tip of the virus. The reference was shifted according to helical symmetry and the alignment parameters were updated at each position by local refinement until the distance of small circle (ii) and the projection of defined point on the z axis was minimal.

---

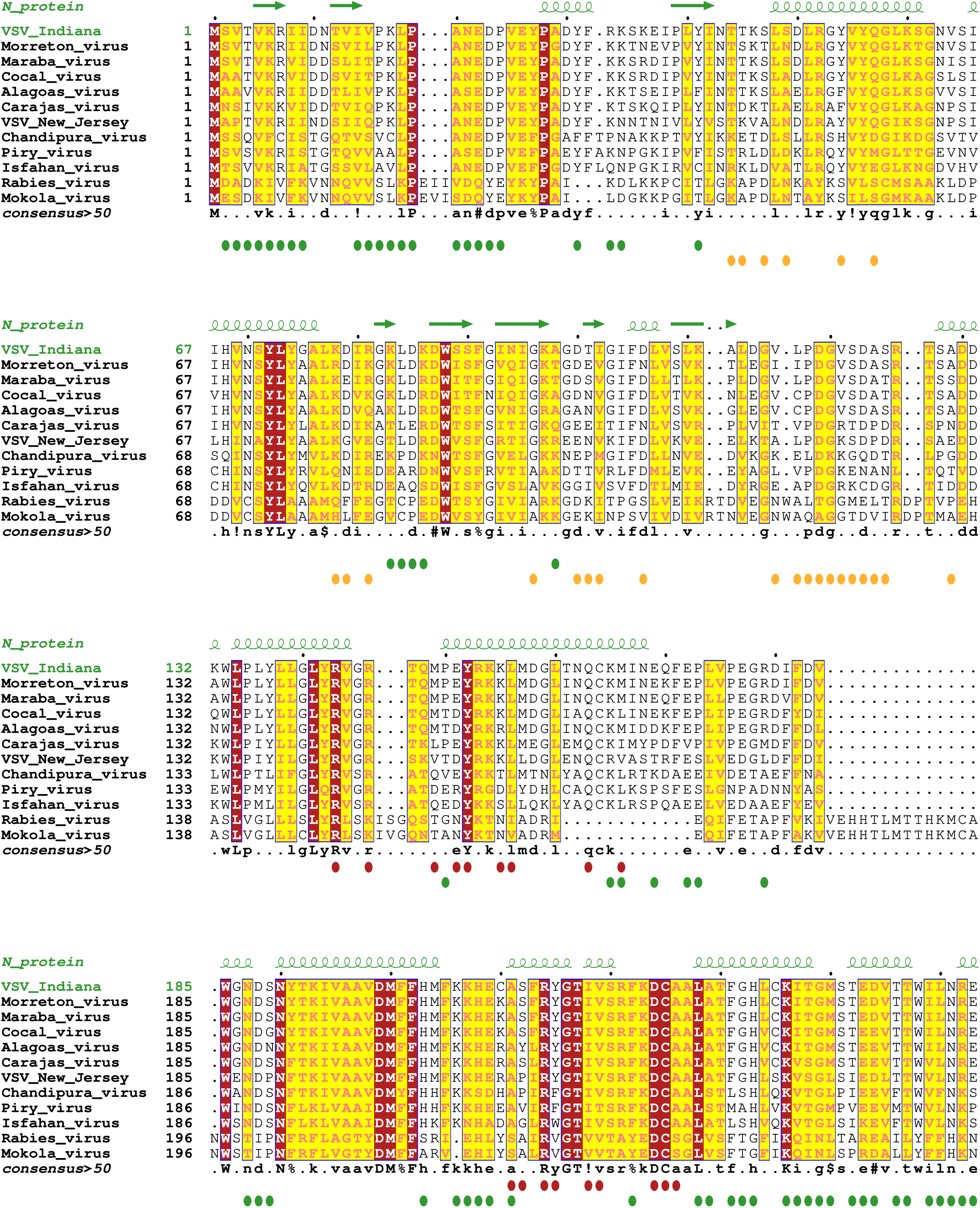
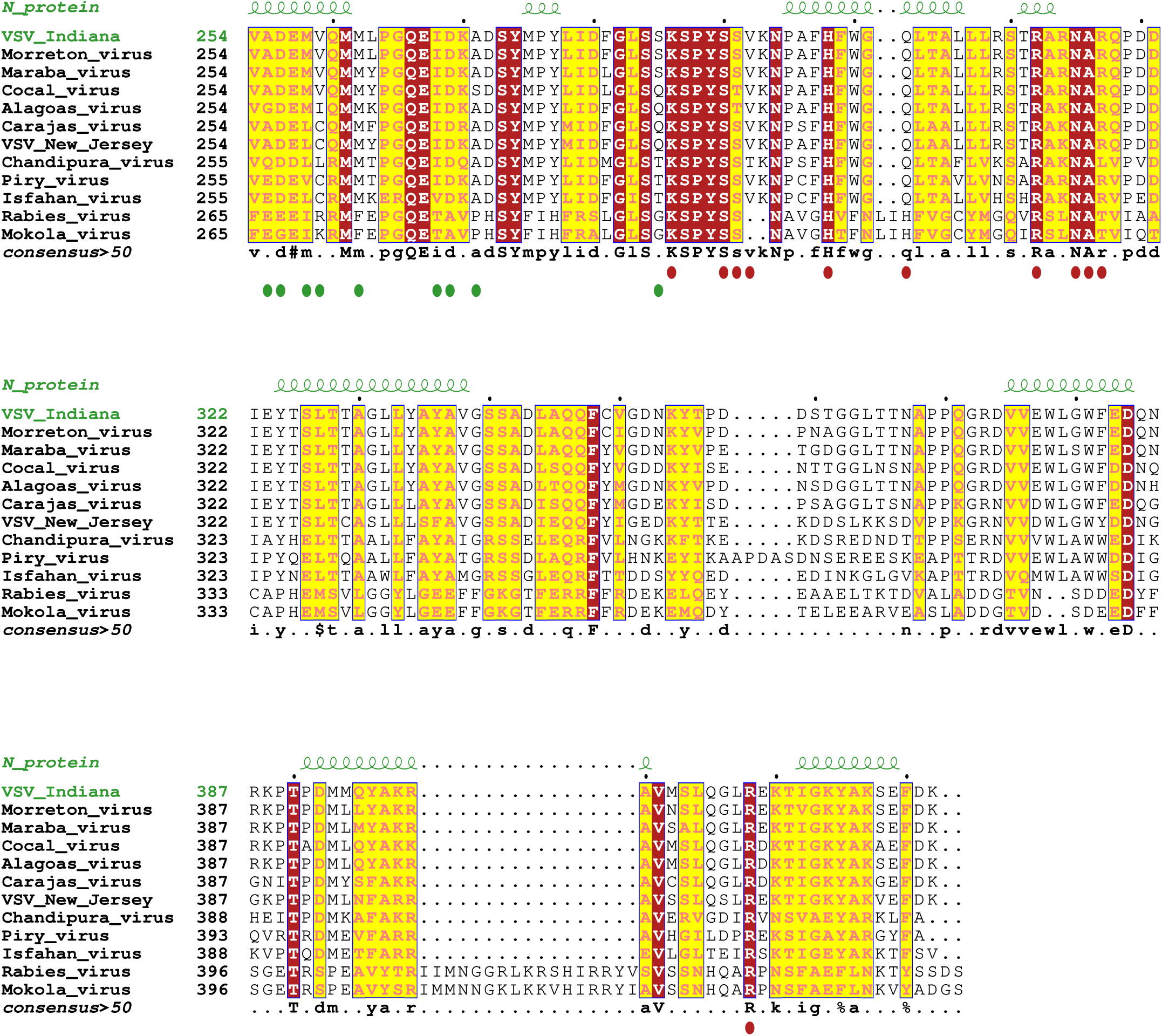
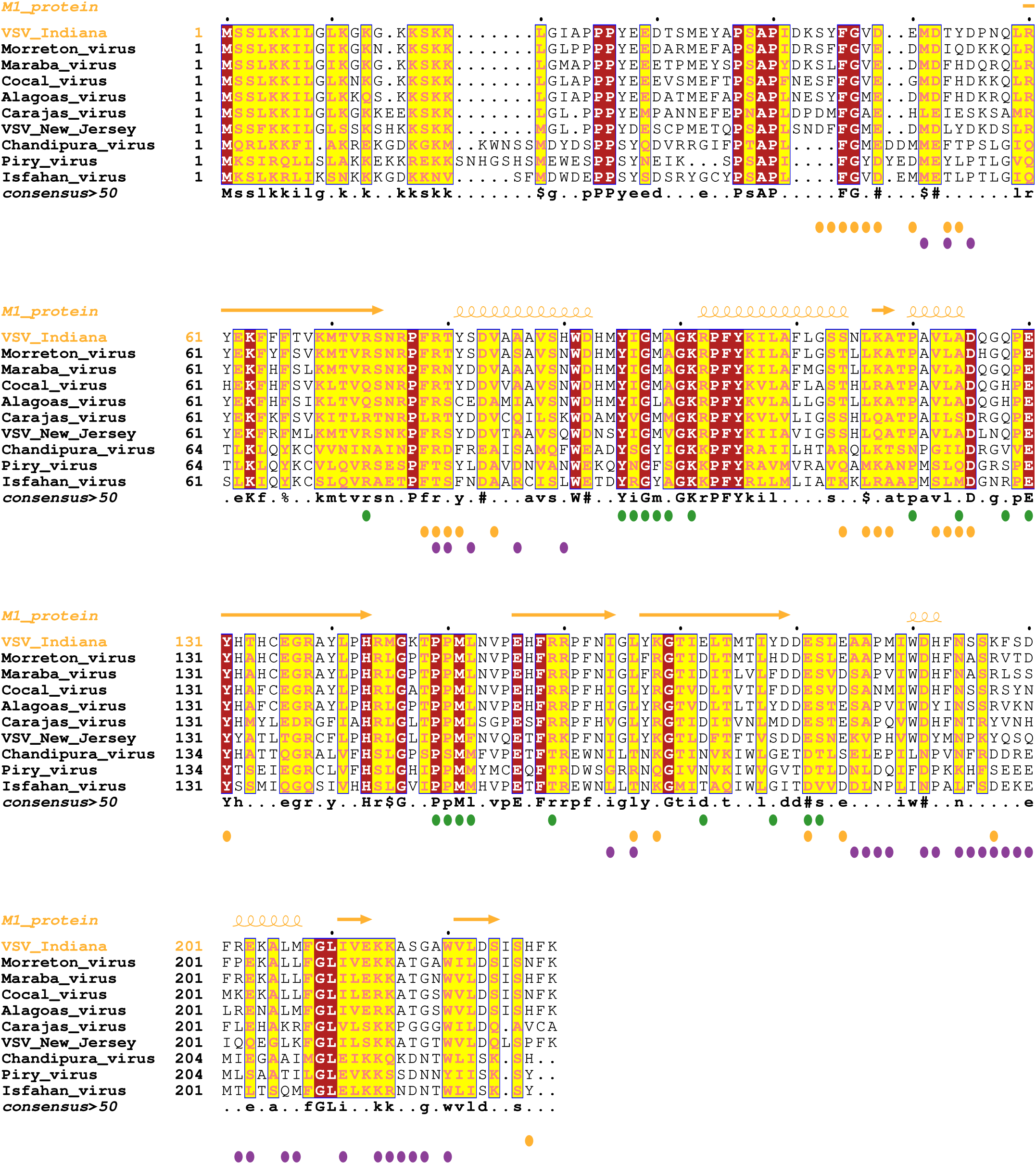
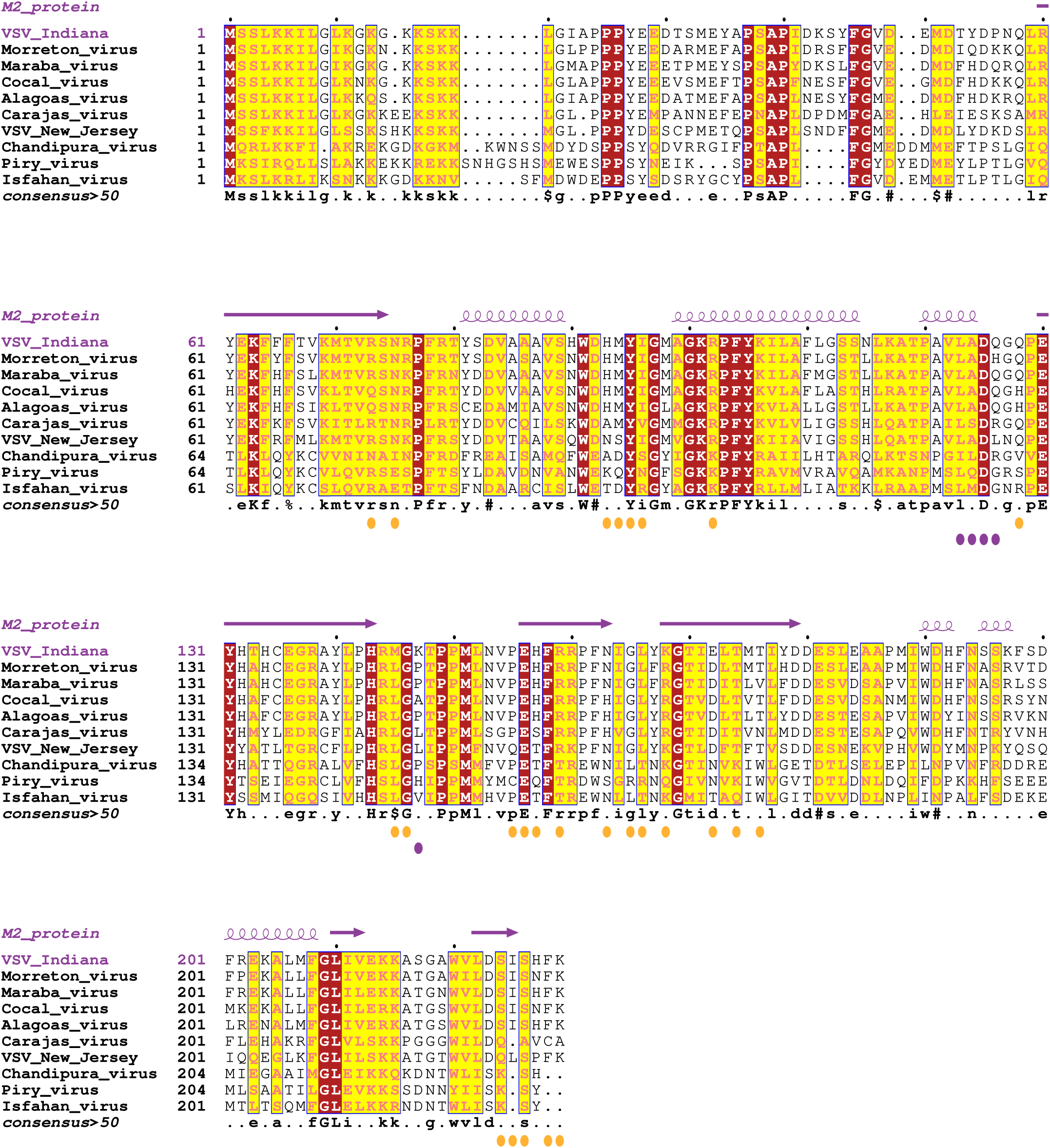

